# Restricted expression site use and extreme genome diversification drives trypanosome antigenic variation in chronic bovine infections

**DOI:** 10.64898/2026.07.31.741217

**Authors:** Stephen D. Larcombe, Jane C. Munday, Guy Oldrieve, Marija Krasilnikova, Craig Lapsley, Craig Duffy, Christina Vrettou, Edith Paxton, Monica R. Mugnier, Andrew P. Jackson, Liam J. Morrison, Richard McCulloch, Keith R. Matthews

## Abstract

*Trypanosoma brucei* exploits an extreme form of antigenic variation to escape the mammalian immune response. This involves the progressive expression of antigenically distinct variant surface glycoproteins (VSGs) on the surface of individual parasites in the population, generating waves of parasitaemia that are successively cleared by host antibodies. Current paradigms were established using in vitro studies and acute rodent infections characterized by high parasitaemia, but natural livestock infections are characterized by low parasitaemia and chronicity. Here, we analysed the infection dynamics of isogenic parasites in mice and cattle in blood during early and chronic infections, quantitating VSG expression diversity within and between hosts, antigen type persistence in vivo and their timing of appearance. This revealed enhanced antigenic diversity in cattle but with a surprisingly reproducible temporal expression hierarchy of related VSGs between independent chronic infections. Analyses demonstrated the unexpected dominance of a single telomeric VSG expression site irrespective of host species and time of infection. Detailed prediction of mosaic VSG assembly reveals exceptional parasite genome diversification within infections involving extensive macro and micro-homology-based recombination to evolve the antigen repertoire. This diversity was restricted but not eliminated in homologous recombination mutants, which could nonetheless sustain chronic infections in mice. These data provide the first comprehensive insight into trypanosome antigenic variation in the clinically-relevant host.

## Introduction

African trypanosomes are parasites responsible for Human and Animal African trypanosomiasis^1,2^, threatening populations and economic prosperity across much of sub-Saharan Africa, where they are transmitted by tsetse flies. Trypanosomes are exclusively extracellular and evade mammalian immune responses by their capacity to change the surface antigen they express in the most extreme form of antigenic variation (AV) identified in a pathogen^3,4^. Individual parasites are uniformly coated by a single type of variant surface glycoprotein (VSG) but can change antigenic type by expressing a new VSG gene at a telomeric VSG expression site. Although only one VSG expression site is active at a time, there are ∼20 VSG bloodstream expression sites (BES) on *Trypanosoma brucei*’s 11 megabase chromosomes and on some submegabase chromosomes ^5,6^. Switches in expressed VSG can involve turning off transcription from the active BES and turning on transcription from one of the other BESs. Additionally, there is a very extensive repertoire of thousands of silent intact VSG genes and pseudogenes housed within subtelomeric gene clusters on megabase chromosomes or at the ends of the parasites’ intermediate and ∼100 minichromosomes. These silent VSGs can be activated by their relocation to an active expression site via gene conversion as an intact gene copy, or by mosaic assembly through segmental gene conversion of VSG genes and pseudogenes^7^. Analysis of *T. brucei* RAD51^8^ and BRCA2^9^ mutants indicates loss of homologous recombination (HR) impairs at least the former reaction, as well as impeding VSG diversification after the induction of a DNA break within the gene ORF^10^, implicating HR in AV. The size of the *T. brucei* VSG repertoire and the combined use of transcription - and recombination - based switching is unprecedented across pathogens that use AV.

Current paradigms of VSG expression dynamics during trypanosome AV are mainly based on in vitro studies, most recently where VSG change is effected through the experimental induction of a DNA double-strand break ^10–12^, or in acute infections in laboratory mice ^13–15^, mostly over the first wave or two of parasitaemia where parasite numbers in the blood often reach more than 10^8^ parasites/ml. Typically, the infecting antigen type dominates until immune clearance of the first peak of parasitaemia, and antigen profiling thereafter indicates a procession of major antigen types, apparently dictated by the genomic location of the VSG donor ^16,17^. However, deep sequencing approaches have revealed that even early in experimental murine infections far more complexity in VSG transcript expression is detectable at the population level, with ∼30- 180 distinct VSG types identified over the first weeks of infection^14,15,18^. As well as the blood, parasite populations are also prevalent in tissues ^19,20^ and there is recent evidence for these sites being less susceptible to immune clearance and important for the generation of new antigenic variants that subsequently appear in the bloodstream^13^. In vitro, DNA double-strand breaks induced within the actively expressed VSG expression site or coding region influence the selection of newly expressed VSGs, both through donor locus choice ^11^ and VSG sequence homology ^10,12^. Upstream sequence homologies between non-VSGs genes in the expression site can also be exploited to effect VSG switches^12,21–23^.

Although African trypanosome AV paradigms have been established in vitro or in mice over 45 years, disease relevant infections are most significant in bovines^24^, with 120 million cattle at risk and 3 million fatalities per year. Here, infections can be sustained for months to years and parasitaemias are considerably lower than in rodents, with parasite numbers in chronic infections typically being close to the microscopy detection threshold ^25,26^. Also, the prevalence of the ‘stumpy’ developmental form that is adapted for transmission to tsetse flies is less than in mice ^26,27^. Whereas the profile of a few dominant VSGs in cattle has been analysed ^16^, the detailed characteristics of VSG expression dynamics in chronic bovine infections are unknown in terms of the scale of population antigenic diversity, hierarchy of antigen expression and influence of donor VSG location or sequence during gene conversion into the expression site(s). Moreover, no work has tested the recombination mechanisms that catalyse AV across long-term infections in any host.

Here, we sought to address these major gaps in our understanding of trypanosome AV in a natural host. By comparison between mouse and bovine infections, we reveal enhanced diversity of antigen expression in cattle and demonstrate a remarkably consistent temporal hierarchy of expression for related VSGs over a considerable length of infection. Also, by exploiting the enhanced resolution enabled by the sampling frequency possible in bovine infections, the dynamics of the appearance and disappearance of antigen types in vivo could be tracked. Notably, genomic interrogation of parasites early and late in infection demonstrated that antigen expression was associated with recombination reactions focused on a single VSG expression site regardless of the host species. Moreover, apparent macro and microhomology based VSG gene diversification by mosaic assembly was prevalent at all infection stages, with a dominant but not exclusive role for RAD51-dependent homologous recombination mechanisms. These findings significantly extend existing understanding of antigen gene activation, expression site usage and VSG diversification.

## Results

### Infection profiles in mice and cattle

To compare *T. brucei* AV in mouse and bovine hosts we established the experimental regimen set out in Figure 1a. *T. brucei* EATRO 1125 AnTat1.1 parasites were inoculated into two groups of eight mice, allowing terminal analysis of blood in the early phase of infection, after clearance of the initiating antigenic variant (peak 2; 8 mice, ‘early’), and at around 30 days, the latest stage of infection ethically permissible (8 mice, ‘late’). For cattle, *T. brucei* EATRO 1125 AnTat1.1 parasites were derived from infections maintained for 60-70 days, with ∼18 samples/animal isolated for transcript profiling, with further samples isolated around day 11 post infection (peak 2) and at the end of the experiment (day 66-70) for parasite cloning and genomic DNA analysis. RNA was analysed by a combination of Illumina and PacBio sequencing and processed for VSG expression analysis via VSGSeq2, a refined deep sequencing analysis pipeline that quantitates expressed VSG transcripts in samples ^28^. RNA samples were also used to monitor levels of PAD1 mRNA ^29,30^ as a marker of stumpy form development at each point in the infections. Genomic analysis of clones was by Oxford Nanopore technologies (ONT), PacBio and Hi-C sequencing methodologies (Supplementary data 1).

**Figure 1.**
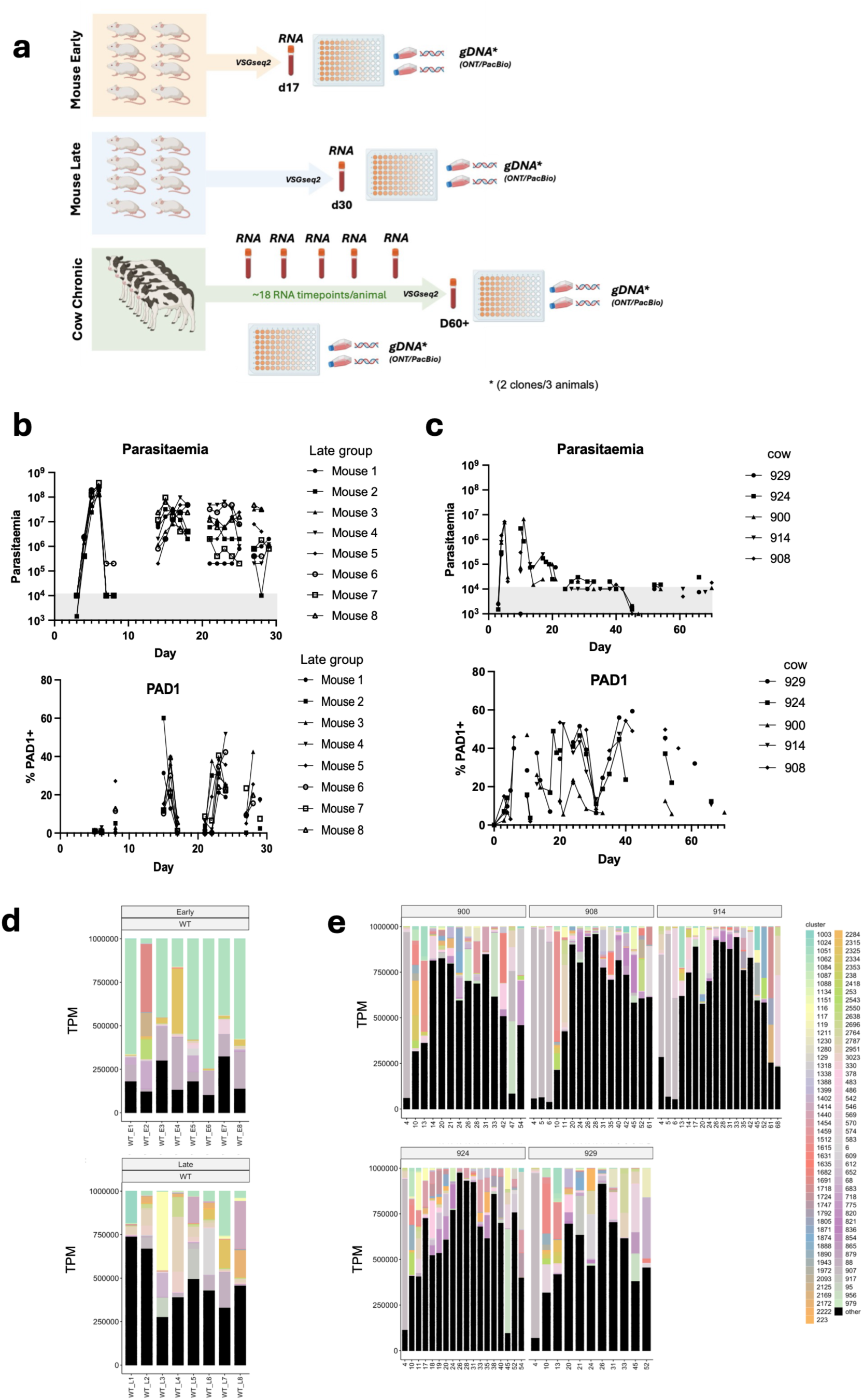
(a) Experimental regimen for the analysis of *T. brucei* EATRO 1125 AnTat1.1 infections in mice and cows. For murine samples RNA was isolated at day 17 (peak 2) and day 30. Cow samples were derived at regular intervals over a 60-70 day infection time course, generating ∼18 samples/animal. Parasites were also isolated and cloned, with genomes being derived by a combination of PacBio, ONP and Illumina sequencing. Genomic sequence was derived from the starting parasites and from 2 clones from each of 3 animals on d11 and d30 (mice) or from d11-17 and d66-70 (cows; animal 908, 914, 924). Images created in Biorender (https://www.biorender.com). (b) Parasitaemias from 8 mouse infections over 30 days; the stumpy marker PAD1 was monitored by qRT PCR and quantitated with respect to a uniform stumpy population (100% PAD1+). The grey zone indicates where parasites were at or below the limit of detection by microscopy (∼10^4^/ml). (c) Parasitaemias from 5 cow infections over 70 days; the stumpy marker PAD1 was monitored by qRT PCR and quantitated with respect to a uniform stumpy population (100% PAD1+). The grey zone indicates where parasites were at or below the limit of detection by microscopy (∼10^4^/ml). (d) Abundance of discrete dominant (i.e. >10% of the overall population) VSG mRNAs in mice, with distinct VSG clusters colour coded. Black bars represent all minor VSGs (>0.01%<10% of the overall VSG mRNA population) combined. WT_E1-8 is mouse 1-8 ‘early’ sample (isolated day 17 post infection); WT_L1-8 is mouse 1-8 ‘late’ sample (isolated day 30 post infection). Y axis= TPM. (e) Abundance of discrete dominant (i.e. >10% of the overall population) VSG mRNAs in different calves (900, 908, 914, 924, 929), with distinct VSG clusters colour coded. Black bars represent all minor VSGs (>0.01%<10% of the overall VSG mRNA population) combined. Day of sample isolation is on the X axis; Y axis= TPM. VSG cluster identities for panel d and e are indicated at the right (Supplementary data 2).

Figure 1b and Figure S1 show the parasitaemia of the chronic infections in mice (‘late’ samples). There was the anticipated major peak of parasitaemia at day 7 post infection (>10^8^ parasites/ml) followed by immune clearance and then recrudescence of parasite numbers by day 13, with parasitaemias then being sustained until day 30. Rapid clearance of parasites in the first peak of parasitaemias in C57BL/6J mice prevented the early dominance of stumpy forms typical in other mouse strains (e.g. MF1 ^30^), but in the chronic phase PAD1 expression was elevated when parasite levels were high (>10^7^ parasites/ml). In cattle (Figure 1c, Figure S1), all animals demonstrated a first peak of parasitaemia on day 4 at ≥10^6^ parasites/ml, with cow 900 showing a second peak at day 10 approaching 10^7^ parasites/ml. After this point all animals sustained infections at approximately 10^4^ parasites/ml, close to the microscopy detection limit. Expression of the stumpy form specific PAD1mRNA remained detectable, supporting the prevalence of parasites expressing some stumpy characteristics in this chronic phase of infection^26^. These infection profiles match previous characteristics of trypanosome infection in mice and cattle^26,30,31^, with an initial peak of parasitaemia followed by a chronic phase, with rodent infections being sustained at parasitaemias around 100-1000 fold higher than those detectable in cattle.

### The diversity and persistence of antigen types in natural infections

To analyse VSG expression profiles throughout infections, RNA from ‘early’ and ‘late’ stage mouse infections, and intermediate RNAs across the duration of the 60-70 day cattle infections were analysed by VSGseq2, with closely related VSGs (>94% identity) being grouped into clusters using cd-hit-est ^28^ to simplify the diversity of highly similar sequences (Supplementary data 2, Supplementary data 3). In ‘early’ mouse infections VSG diversity was relatively consistent in 6/8 animals, with the same dominant VSG clusters accounting for 60-80% of transcripts, whereas up to 20% of assignments were to minor VSG transcripts (i.e. those representing 0.01% - 10% of the overall population) (Figure 1d). This matches the diversity previously observed in rodent infections by deep sequencing approaches ^14,15^; indeed, total numbers of VSG clusters detected (∼100-200; Figure 2a) were comparable to previous values at an equivalent infection stage ^15^. In cattle, Cluster 907 (which represents VSG AnTat1.1 expressed by the inoculated parasites) dominated in all animals on day 4 but from day 13-17 (peak 2) and beyond, minor VSGs dominated (Figure 1e). By day 28-30, overall VSG diversity increased in mice and cattle, with 25 - 60% of VSG clusters detected comprising minor VSGs in mice and >80% in cattle, with over 200-400 detected VSG clusters expressed by at least 0.01% of the overall population (Figure 2a). Thus, despite the much lower parasitaemia in calves, VSG diversity arose more quickly and with greater complexity (Figure 2a). After 30 days in cattle, the overall number of VSG clusters mapping to the whole VSGome decreased in all animals, although sampling frequency and the recovered parasite number decreased from this point particularly late in infection when abundant bovine material confounded parasite isolation.

**Figure 2.**
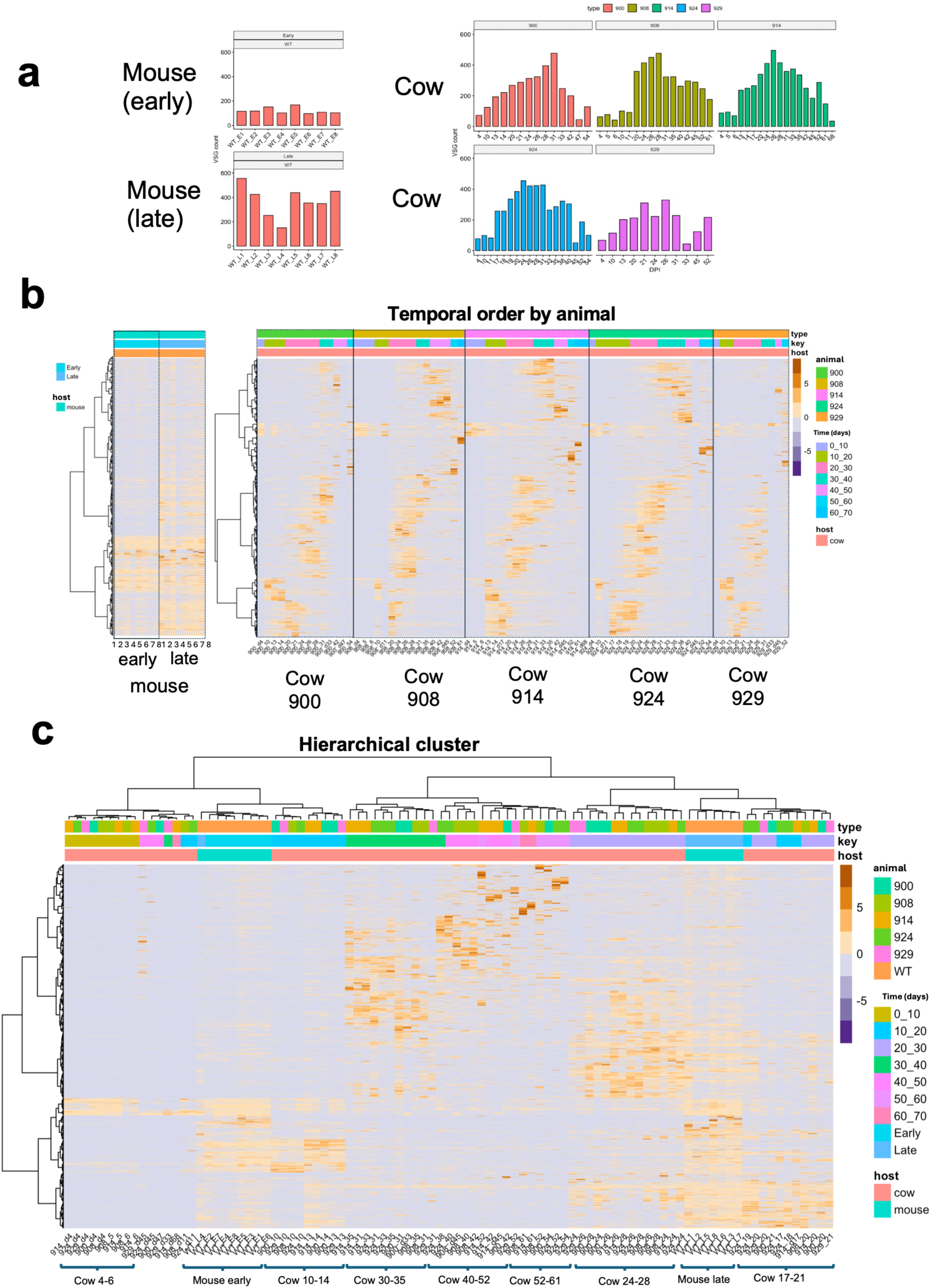
(a) Expressed population diversity of VSG clusters in mice (8 early and 8 late time point) and in 5 calves over d4 to d54-68 days of infection (at >0.01% of the population total) (b) VSG clusters expressed in 8 early mouse samples and 8 later mouse samples, and in 5 cows over the time course of chronic infections, arranged hierarchically within each group (early, late mice) or individual animal (cows) (c) VSG clusters expressed in 8 early mouse samples and 8 later mouse samples, and in 5 cows over the time course of chronic infections ordered by hierarchical clustering of relatedness indicated on the left hand dendrogram.

Figure 2b and Figure S2a shows VSG clusters ordered by host animal and time, revealing striking consistency in the expressed VSG profiles between infections. Moreover, by grouping samples hierarchically by relatedness (Figure 2c), the temporal expression profile of VSG clusters was found to be similar between mice and cows, with early mouse samples (d17) grouping with cow samples around d10-d14, and late mouse samples (d28-30) grouping with cow samples at d17-28. Despite this, comparison of the expression of individual, rather than closely related, VSG transcript sequences between mice and cattle across the course of their infections demonstrated that only 5.0% of VSGs were expressed in both hosts (at any timepoint), with 88.5% being expressed only in cattle and 6.5% expressed only in mice, re-emphasising the enhanced diversity of antigen expression in bovine infections (Figure S2b). In summary, antigen diversity was higher in cattle than mice and clusters of related VSGs were expressed in a consistent temporal profile between animals, supporting an underlying hierarchy of expression. Nonetheless, distinct individual VSGs were expressed in each animal at each stage of infection.

The ability to monitor the temporal expression of VSGs in individual cows allowed some facets of VSG dynamics to be quantitated. For example, to compare the persistence in blood of parasites expressing particular VSG clusters, dominant VSG clusters (i.e. representing >10% of the VSG population) were aligned with respect to their first appearance and subsequent disappearance in consecutive samples, revealing that VSG clusters were reproducibly detectable in blood for 5 days regardless of their time of peak abundance during the infection (Figure 3a, left). Although some low-abundance VSG clusters persisted longer, most persisted for 5 days regardless of their prevalence in the population (Figure 3a, middle) and irrespective of VSG protein length (Figure 3a, right), which also did not affect the timing of expression of VSGs (Figure S3). Hence, closely related VSGs appear to have a quite consistent lifespan in the blood, with later expressed and minor VSGs not escaping immune detection and clearance despite evidence of infection-related pathology, reflected by a reduction of haematocrit in all animals beyond day 20 (Figure S4).

**Figure 3.**
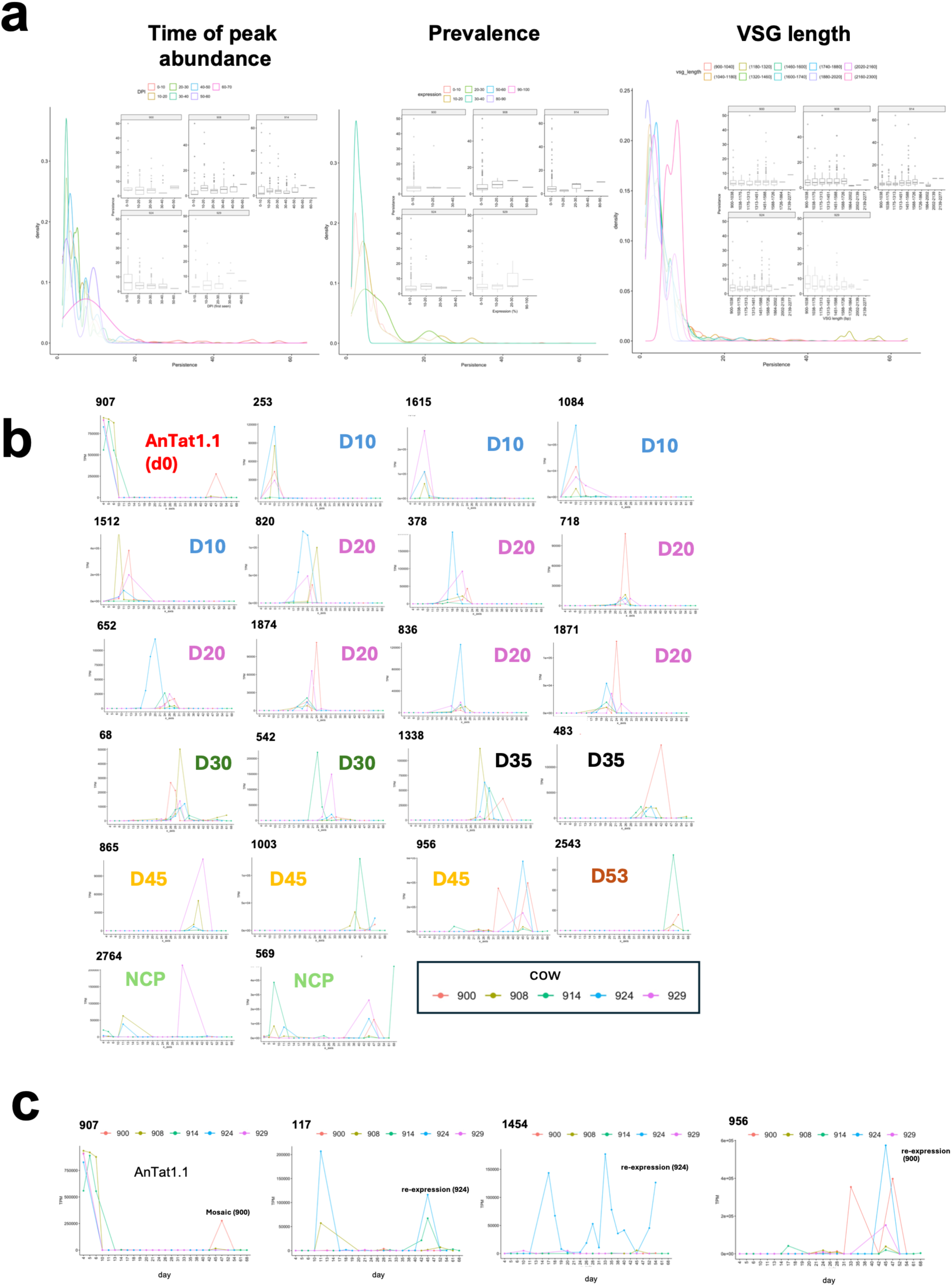
(a) left; Persistence of VSGs (time from first appearance to last appearance in consecutive samples) that show their peak abundance at different timepoints during cattle infection. Throughout the time-course of infections the persistence of VSGs is around 5 days; *middle;* persistence of VSGs (time from first appearance to last appearance in consecutive samples) whose abundance is in each percentile of the overall prevalence in the population. Most VSG groups persist equivalently although some minor VSGs are detectable longer. *right*; persistence of VSGs (time from first appearance to last appearance in consecutive samples) plotted for VSGs of different lengths. In each case, the main plot combines data from all animals; data from individual cows is inset. (b) Expression profiles for individual VSG clusters that are expressed in 4 or more calves over time. Each title (D10, D20 D30, etc.) represents the approximate peak of expression, NCP= no consistent profile of expression between animals. (c) Expression profiles for individual VSG clusters that are re-expressed during the course of infection. Where VSG clusters are re-expressed later in infection, individual transcripts from individual animals at the time point where a VSG cluster was re-expressed were examined for evidence of being mosaic or directly re-expressed as an identical sequence to the earlier infection timepoint.

### Remarkable temporal hierarchies in VSG expression between animals

To examine the temporal hierarchy of VSG cluster expression between individual animals, the VSGs expressed in the five bovine infections were compared. Most (63%) individual VSGs were specific to an individual animal (408/645 VSGs; Figure S2c) but 22 dominant VSG clusters (i.e. comprising >10% of the total population of expressed VSGs) were detected in at least 4 animals, allowing their temporal expression profile to be compared (Figure 3b). Strikingly, consistency in the temporal expression of these VSG clusters was seen at each phase of the infection in each animal, despite differences in the relative expression abundance of clusters between animals. Figure 3b shows VSG clusters dominant at approximately d10 (e.g. VSG 253), d20 (e.g. VSG820), d30 (e.g. VSG68), d35 (VSG 1338), d45 (e.g. VSG 865) and d53 (e.g. VSG 2543), whereas some clusters showed no consistent expression profile between animals (NCP; e.g. VSG 2764, 569). Extending the analysis to the 94 dominant VSG clusters shared in 3 or more animals further supported a structured temporal hierarchy extending throughout the chronic cattle infections, with reproducible expression profiles of related VSGs at equivalent stages of infection (Figure S5). Analysis of the temporal profiles of VSG expression also provided evidence of the reappearance of some VSG clusters, as either a mosaic or direct reexpression of the same VSG. For example, cluster 907 (representing the initiating VSG, AnTat1.1) was expressed by all animals at day 5, but in one animal (900) a cluster 907 VSG sequence was also detected toward the end of infection as an apparent mosaic (Figure 3c; Figure S6a). However, a number of other related VSG cluster sequences showed evidence of direct re-expression of individual VSGs without mosaic formation (VSG 117 and VSG1454 in cow 924, VSG 956 in cow 900) (Figure 3c, Figure S6b-d), indicating that some VSGs can re-emerge after clearance earlier in the infection.

### The genomic context of expressed VSG in the starting genome

The genomic context of expressed VSG in mice and cows was assessed by assembling genome of the *T. brucei* EATRO 1125 AnTat1.1 parasites used to initiate infections, and then mapping expressed VSGs at different points of infection. PacBio, ONT and Hi-C were used in combination to generate a high-quality genome assembly, with HiFiasm^32^ and Verkko^33^ used to assemble the megabase genome, retaining the diploid core genome, with haploid subtelomeres and VSG expression sites resolved using Flye^34^ (Figure S7). Of 41 chromosome ends identified, 18 were assigned as BES based on their possession of promoter sequences, expression site associated genes and 50bp and 70bp repeat regions (Figure 4a), and a further 7 were assigned as metacyclic ES based on the presence of previously identified M-VSGs and an absence of BES features ^35^. The AnTat1.1 VSG gene was present as a subtelomeric silent gene inversely oriented with respect to the Chromosome 8 telomere, and its upstream co- transposed sequence^36^ was detected in an expression site on telomere 1iiB although the telomere 1iiB sequence did not fully extend to the VSG and telomere end. However, the presence of the AnTat1.1 gene at this location was consistent with its ubiquitous expression in parasites used to initiate the mouse and cow infections and with earlier expression site mapping^37^.

**Figure 4.**
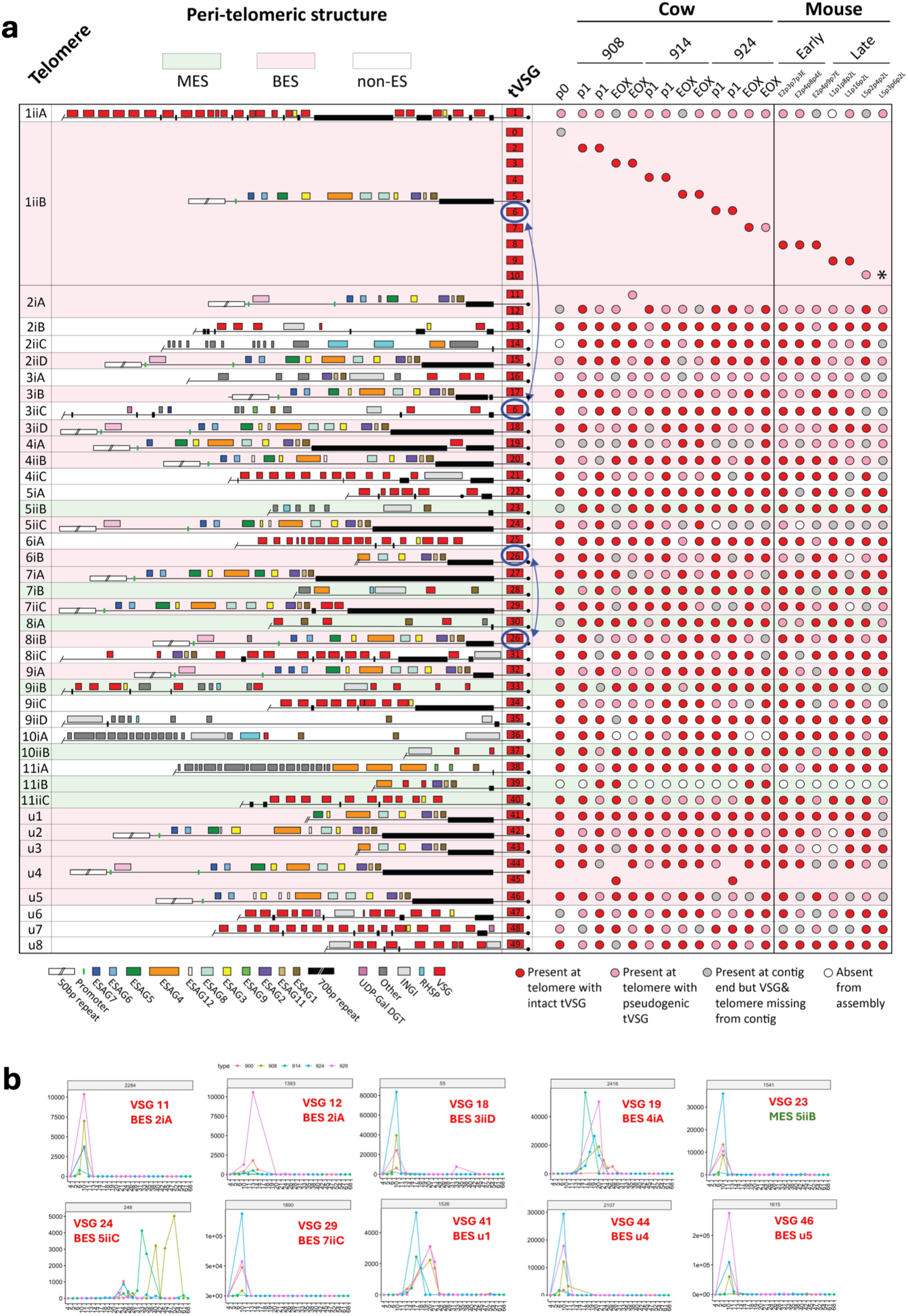
(a) Map of the telomeric EATRO 1125 bloodstream (BES) and metacyclic (MES) expression sites in the starting genome. ES elements are colour labelled according to the key at the bottom of the map. Peri-telomeric sequences are ordered and named according to chromosome. The observed chromosome number (1-11) is assigned where identifiable in these clones, followed by haplotype (i or ii) and then a letter identifier (A-D). Eight peri-telomeric sequences that belonged to unassigned contigs that could not be associated with a chromosome are labelled u1-u8. VSG genes found in each EATRO 1125 BES are annotated for each clone analysed, derived either at the initiation of infection (p0), early (p1) or end of experiment (EOX) for the infection for cow derived clones, or clones derived from mice. Cows from which each clone were derived are 908, 914 or 924; mouse clones were derived from either the early or late infection group. Different telomere proximal VSGs (tVSGs) are numbered (0-49) and coloured coded to reflect them being likely intact VSGs (red) or VSG pseudogenes (pink). Blue arrows indicate two cases where the same tVSG sequence is found in different positions (i.e. tVSG6 in 1iiB/3iiC and tVSG26 in 6iB/8iiB). The AnTat1.1 gene is inferred to be at the p0 position on telomere 1iiB based on expression and sequence of the co-transposed segment at that location. In clone L5p2p4p2L, the VSG locus is not assembled and cannot be inferred (*). (b) Expression profile of tVSG genes located in other telomeres distinct from EATRO 1125 BES1 in each animal through the duration of infection. The tVSG identifier and telomeric location is indicated in each case. The predicted metacyclic expression site MES 5iiB (green) harbouring VSG23 is indicated.

To explore the genomic context of VSGs as they are activated during infections, parasite clones were isolated from mouse infections (early and late) and from d11 or d17 (peak 2) or d66-70 (end of experiment, EOX) of bovine infections. For cattle- derived samples, pooled parasite populations were established from the blood of each of three cows (900, 908, 924), this requiring culture adaptation, and two clonal populations were derived from each pool. RNAseq analysis established that the derived clonal populations each expressed a single major VSG, which was representative of the dominant VSG expressed in each established pooled population prior to cloning (EOX clones are shown in Figure S8a). Overall, the genomes of the 19 clones from mice and cattle were sequenced to determine VSG gene occupancy at telomeric expression sites in each (Figure 4a, Figure S8b). Surprisingly, changes in VSG gene ES occupancy in each clone from each animal were restricted to the same BES used in the starting population at telomere 1iiB (hereafter named EATRO 1125 BES1). Thus, in clones derived from different animals isolated at different times from either cattle or mice, different VSGs occupied EATRO1125 BES1, contrasting with other VSG expression sites, where the same telomeric VSG (tVSG; either intact or as a pseudogene) persisted at the same location throughout the course of infection (Figure 4a). The *in vivo* relevance of the VSG occupying EATRO1125 BES1 in gDNA was confirmed by the expression, in multiple animals, of VSG clusters representing tVSGs 4, 6, 8 and 9 in both mice and cows, at appropriate timepoints in the infection (e.g. clusters 570, 683, 1414 and 2284 Figure 1e; Figure S5). Interestingly, although the expression of VSGs from other telomeres was not detected in any of the clones, expression of other tVSGs was detected by VSGSeq2 at a population level early in infection (usually prior to day 20; Figure 4b), with a temporally similar peak expression profile in distinct animals. This provided evidence that other BES-resident VSGs were either activated in situ soon after clearance of the initiating AnTat1.1 VSG, with a similar hierarchy of activation and expression between animals, or they were recombined into EATRO1125 BES1 for expression at this timepoint. Evidence for the latter situation was evident in a cow 924 day 11 derived clone, where the VSG gene on telomere 3iiC (VSG 6) was copied into EATRO1125 BES1 for expression (Figure 4a), and some other expressed telomeric VSGs resided at sites lacking BES features (e.g. VSG 23 was resident in a predicted metacyclic ES). Although two other BES showed a change in the resident VSG (telomere 2iA in cow 908 clone EOX2, and telomere u4 clone p1 in cow 908 and 924), these VSGs were not detected as dominant transcripts by VSGsSeq2. Thus, throughout infection in mice and cattle, EATRO1125 BES1 was almost exclusively responsible for VSG expression and for antigen switching.

### Extreme antigen gene diversification throughout infections

In addition to examining the contribution of the expression site used during antigen switching, the contribution of mosaic genes was examined. The 28,876 VSG transcripts identified across all cow samples were filtered to select those represented by at least 100x coverage and present in two or more samples to provide the highest possible sequence confidence. These analyses focused on individual transcripts rather than related VSG clusters to ensure the high stringency sequence similarity necessary to identify parent genes and gene fragments in the genome. The high confidence transcripts were then searched against all available EATRO 1125 VSGomes (i.e. a combination of our genome assemblies and all publicly available EATRO 1125 genomes and VSGomes) to provide comprehensive sequence coverage, thereby maximising our ability to identify intact gene copies and putative mosaic donors. Of the filtered subset of 5,889 VSG transcript types, 188 had a >99% match over >99% of the sequence length, indicating that 3.2% of the high confidence VSG transcripts across chronic infections were identifiable as being derived from an intact functional gene. To search for potential mosaic donors for the remainder of this transcript set, high resolution mapping for each transcript was carried out by searching blocks of 50 nucleotides across the transcript length, with a 1bp sliding window. Each segment was aligned to the combined EATRO 1125 VSGomes, also segmented into 50 nucleotide blocks with a 1bp sliding window, with potential mosaic donor regions being identified based on having >99% identity to the VSG transcript. Potential donor sequences could be mapped along 95%±6.6% of the length of members of the 5,889 VSG transcript set (Figure S9). Among the high confidence set, a mean of 1.82±1.17 donors contributed per mosaic, with most contributing 1 donor element with a modal donor fragment length of 104bp (and a mean of 489bp) with 99% coverage of the transcript (a similar profile was observed with a less restrictive dataset allowing >80% transcript coverage, Figure S10, S11). These recombination events were distributed across the coding sequence, but with a notable 3’end bias (Figure 5a). Extending the analysis to all 28,876 expressed VSG transcripts demonstrated a similar profile of potential mosaic donors, with only 2,407 (8.3%) transcripts present as an intact donor gene in any EATRO 1125 VSGome with >99% coverage and with a similar number and distribution of mosaic donors as seen in the high confidence transcript set (Figure 5a, lower). Significantly, the complexity of mosaic formation increased with time, with simple mosaics comprised of only 2 putative donors dominant at day 1-9, whereas at day 50-59, 60% of expressed VSG were assembled from 4 or more putative donors and some had >8 potential donors (Figure 5b). Analysing individual transcripts revealed considerable complexity, with unique or multiple potential donors located across different chromosomes (Figure 5c shows examples of a simple and complex mosaic from day 14 (cow 900) or day 52 (cow 929) (full transcript dataset available at doi.org/10.5281/zenodo.21294599). Among all VSG transcripts in our analysis, 56 had no mappable donor sequence identifiable in the combined EATRO 1125 VSGome (Figure S9a), similar to previous analyses of the *T. b. gambiense* VSGome^38^. Overall, this analysis demonstrates that the VSG gene archive exhibits extreme diversification during chronic infections, with a dominance of complex mosaics assembled from identifiable potential donors that accumulates over time.

**Figure 5.**
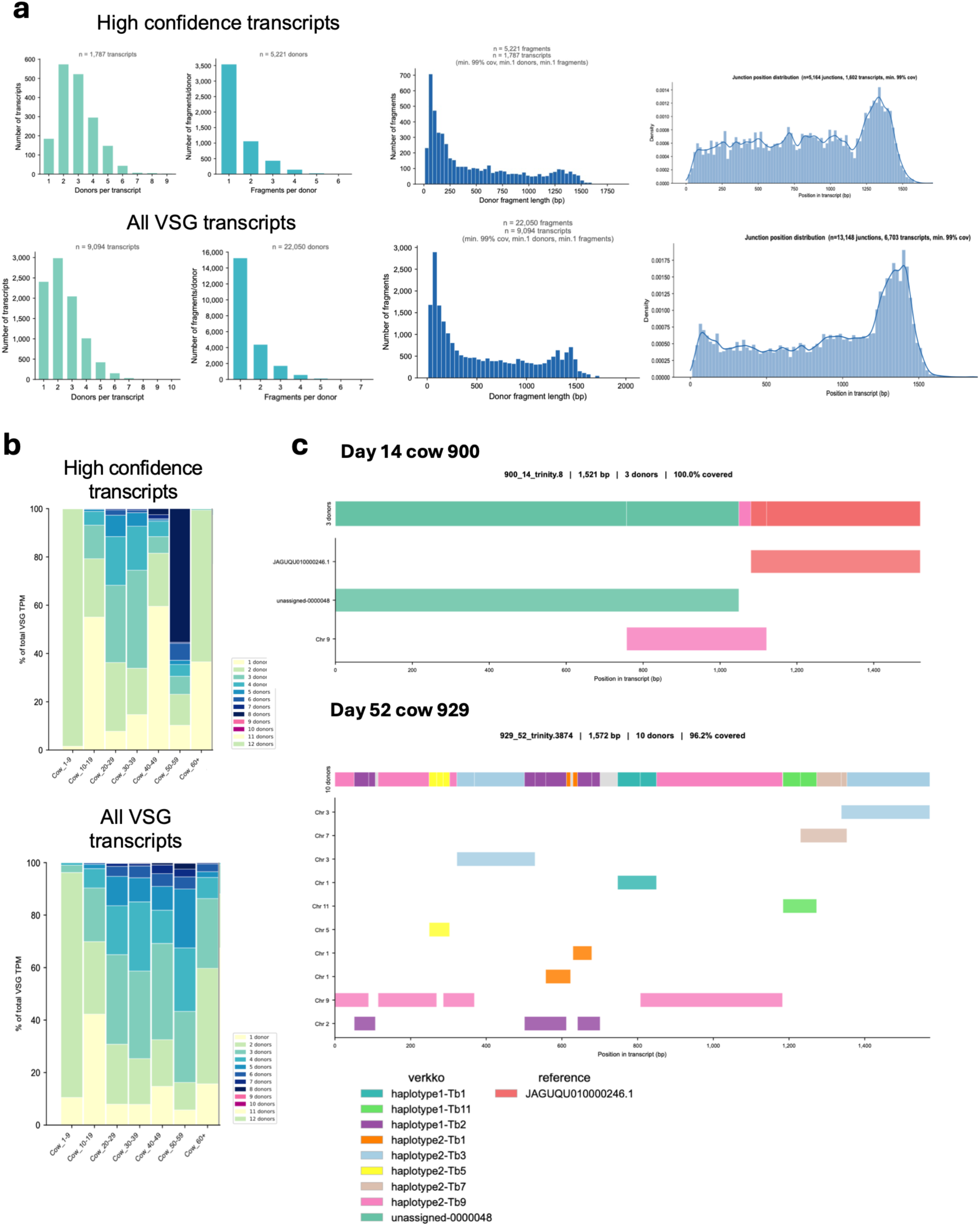
(a) Mosaic formation in the ‘high confidence’ and ‘all VSG’ transcript data set for transcripts where 99% of the sequence is covered by identifiable donors. Histograms show the number of donors per transcript, the number of homologous sequences contributed by a given donor, and the length distribution of donor fragments. Finally, the position of donor junctions is shown across the VSG sequence length highlighting the 3’ end bias of the recombination events. (b) Representation of the number of mosaic donors identified that cover at least 99% of the VSG transcript at different times of infection in cattle for the high confidence VSG transcripts and the entire VSG transcript dataset. Mosaic donors increase with time reflecting increasing mosaic complexity during infections. (c) Representative mosaic derived transcripts highlighting putative donor sequence locations, with regions mapping to one donor or multiple donors being indicated. The upper transcript is derived from day14 of infection for cow 900 and is composed of 3 putative donors. The lower transcript is from cow 929 at day 52 and has 10 putative donors able to provide 100% coverage of the expressed transcript.

### Recombination mutants show reduced VSG diversity but sustain chronic infections

In laboratory adapted *T. brucei* Lister 427, RAD51 and BRCA2 contribute to VSG switching via homologous recombination^39,40^. To explore the role of known homologous recombination mechanisms during VSG switching over long-term infections and in the extreme diversification of the VSG gene repertoire we observed, null mutants for each gene were generated in pleomorphic *T. brucei* EATRO 1125 (Figure S12a-c). Both lines grew more slowly than wild type cells in vitro (Figure S12d), whereas in mice, each mutant generated a first peak of parasitaemia of >10^8^ parasites/ml - equivalent to wild type cells - but thereafter sustained only fleeting infections. This was more pronounced in the RAD51 mutants (∼10-100 fold fewer than wild type) than BRCA2 mutant parasites (Figure 6a) but, nonetheless, infections were sustained over 30 days for each mutant. The VSG expression profile of each mutant was assayed from terminal sampling at the second peak (d15, ‘early’) or after 28-41 days (‘late’) and at the late timepoint parasite clones were derived for genome analysis. At the early timepoint, the diversity of expressed dominant VSG clusters in the mutants was less than in wild type cells, and there were fewer minor (<10% of total) VSG clusters (Figure 6b). At d30, when the proportion of minor VSGs in wild type parasites had increased to approximately 40%, in both RAD51 and BRCA2 mutants these minor VSGs accounted for only 5-10%, with expression of a few major VSGs dominating. These data indicate that loss of either homologous recombination factor impairs the capacity to generate expressed VSG diversity. Hierarchical clustering of the expressed VSGs in the mutants against both wild type mouse and chronic bovine infections demonstrated that the VSG expression profile of the HR mutants was highly restricted compared to wild type cells, whether derived early or late in infection, being most similar to the very earliest (peak 1) bovine samples (d4-10) and dissimilar to the wild type mouse VSG profiles (Figure 6c). As with cow infections, mosaic complexity in wild type parasites in mice increased between early and late infection, such that putative VSG gene donors increased from 1-2 in the early infection group (70% of transcripts) to up to 5 donors in the late group (Figure 6d). In contrast, the frequency of predicted mosaic VSG gene assembly was reduced in the recombination mutants late in infection although the ability to generate potential mosaics in both RAD51 and BRCA2 mutants remained, albeit restricted to the longer donor lengths more typical of early infection in wild type parasites (Figure 6e). Genome analysis of parasite clones from the HR mutants grown in mice further demonstrated that mosaic donors were long and were not restricted to expression sites, with donor sequences distributed across the genome (Figure S13). Interestingly, unlike the parental parasites, the recombination mutant clones exhibited loss of genomic subtelomeric regions, potentially reducing their VSG archive (Figure S14) and overall fitness (Figure 6a, FigS12d). Hence, the establishment of chronic murine infections remains possible in HR mutants and mosaic formation using diverse donors can still be observed but the extent of diversification in each recombination mutant is greatly reduced and this is further contributed to by subtelomeric deletion.

**Figure 6.**
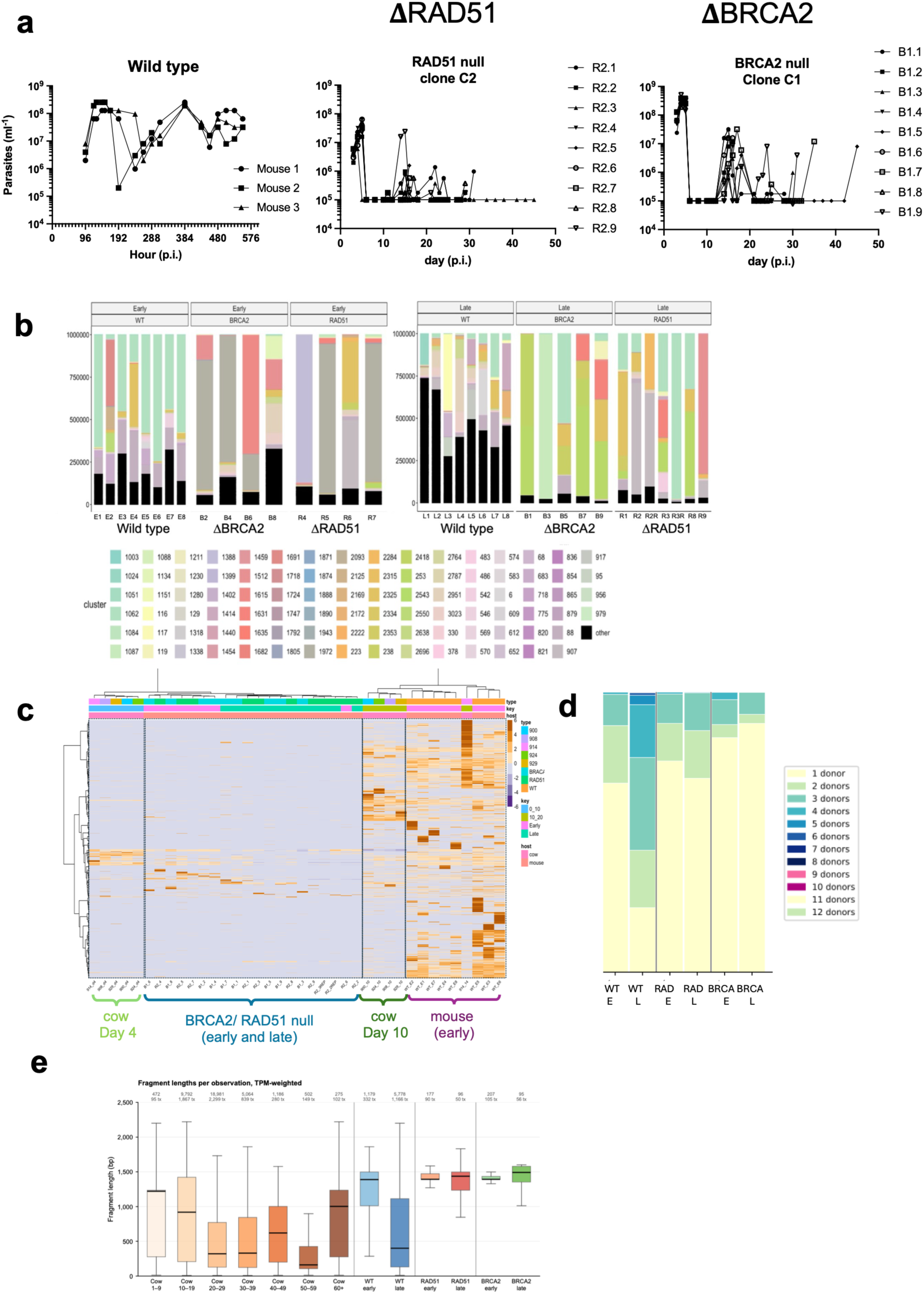
(a) Left; parasitaemia in mice of wild type EATRO1125 AnTat1.1. *Middle* and *right*; parasitaemia in mice of null mutants for RAD51 and BRCA2. After a first peak of parasitaemia each recombination mutant was sustained in mice but at very low parasitaemia with occasional peaks. (b) Abundance of discrete dominant (i.e. >10% of the overall population) VSG clusters in mice, with distinct VSG clusters being individually colour coded according to the key (Supplementary data 2). Black bars represent all minor VSGs (>0.01%<10% of the overall VSG mRNA population) combined. Mice were analysed for each of the RAD51 and BRCA2 null mutants either from day 17 (early) or day 30 (late); wild type parasites (8 mice early, 8 mice late) are those also represented in Figure 1 and 2. Y axis= TPM. (c) Hierarchical ordering of VSG relatedness in mouse (wild type, RAD51 and BRCA2 mutants) and cow samples derived on day 4 and day 10. The recombination mutants cluster optimally with the day 4 cow samples. (d) Representation of the number of mosaic donors identified that cover the VSG transcript (with 99% coverage) at different times of infection (E, early; L, late) for either wild type parasites or RAD51 or BRCA2 null mutants. (e) Distribution of mosaic donor fragment lengths in samples derived from cow infections at different times points, from early and late mouse infections and from the recombination mutants late in infection. The donor fragment length in the recombination mutants was significantly longer than in the wild type parasites in cows or mice later in infection.

## Discussion

Trypanosomes are an established exemplar of antigenic variation as a pathogen immune escape mechanism. However, for almost all studies to date, the analysis of parameters important for antigenic variation have been restricted to in vitro studies with laboratory adapted monomorphic parasites (e.g. ^11,12^) or in vivo studies in non-natural rodent hosts limited to the first weeks of infection (e.g. ^13,14^). This contrasts with natural infections in the field, where infections can persist for months, involving developmentally competent pleomorphic parasites adapted for chronic infections and tsetse transmission, with bovines being the most clinically-relevant and economically important hosts^24^.

In this study we have comprehensively examined, for the first time, trypanosome antigenic diversity in the natural bovine host, with direct comparison to isogenic infections in the murine model. The capacity to analyse longitudinal parasite material with significant temporal granularity has allowed the dynamics of antigen expression to be tracked in distinct infections over time and, importantly, enabled the correlation of these dynamics with genome organisation of the very large repertoire of VSGs. Our analysis has revealed patterns of gene activation for related VSGs that generate broadly conserved hierarchies of antigenic expression in different host species and in independent infections in individuals of the same host. We also find evidence of exceptional VSG gene repertoire diversification throughout infections, which is reduced when major mechanisms of homologous recombination are disrupted.

Several key observations emerge that address long standing questions and existing paradigms concerning trypanosome AV. By tracking the appearance and disappearance of VSGs, the persistence (first detection to disappearance) of individual antigen types in blood was found to be approximately five days, regardless of their abundance or time of appearance during the infection. Thus, dominant VSGs are not preferentially eliminated by the immune system and VSGs do not ordinarily escape detection, although some can persist or be re-expressed later in infection. Moreover, the limited persistence of expressed VSGs in the blood was maintained, even later in infection, when pathology is observed and immune suppression predicted ^41^. Also, we found no evidence for VSG length alone dictating the highly reproducible timing of VSG expression, contrasting with some reports ^11,42^.

Analysis of the VSGs expressed in mice and cattle demonstrated a similar overall hierarchy of expression of related VSGs over time. In particular, hierarchical clustering revealed best correspondence between mice infections on d17 and cow infections on day 10, coinciding with second peak of infection in each host. Similarly, mouse samples derived on d30 best matched cow material harvested on d17-21, emphasising that the elevated diversity of VSG expression in cattle is associated with a more rapid progression through their VSG repertoire. This is perhaps related to the enhanced carrying capacity of these hosts ^43^, with an overall parasite load predicted to be approximately 2.5-25 times larger in cattle than mice accounting for the relative host size (25000x larger for cattle) and parasitaemia (1,000-10,000x higher in mice). As well as between host species, there was hierarchical conservation between animals, with related VSGs between infections exhibiting strikingly similar temporal expression profiles. These findings substantially extend our understanding of the patterning of VSG expression dynamics, which hitherto was limited to the relatively predictable VSG expression profiles detected in the early stages of *T. brucei* infection in rodents^44,45^. The findings also have parallels with AV in other *Trypanosoma* species, since order of VSG expression during long term *T. vivax* and *T.b. equiperdum* infections has been documented in goats and rabbits, respectively ^46–48^.

Contrasting with these other trypanosome species, only in *T. brucei* do we have a clear understanding of the organisation of the VSG repertoire in the genome. A smaller scale earlier analysis of *T. brucei* VSG expression timing in cattle infections ^16^ suggested the temporal appearance of VSGs corresponded to the genomic position of each antigen gene at the outset of the infection, whereas in mice VSG length has been proposed to contribute to expression priority ^42^. Our data instead supports sequence relatedness with the previously expressed VSG or its surrounding sequences as the most important determinant of expression hierarchy. This is consistent with the concept predicted by modelling ^17^ of the expression of ‘strings’ of VSGs related by sequence and extends it to much earlier stages of infection than previously anticipated. Although our analysis focused only on the bloodstream parasite population due to the intractability of sampling bovine tissue material at scale (and in particular the difficulty in perfusing cattle tissues/organs in order to remove the confounding circulating blood), recent studies in mice have demonstrated that the blood VSG profile reflects the overall tissue VSG population albeit with less complexity, possibly due to enhanced rates of VSG clearance in the vasculature ^13^.

To better understand the relative contribution of in situ activation by transcriptional switching between VSG expression sites and recombination of VSGs into expression sites, we derived high quality genomes of the infecting population and 19 clones of parasites derived from different animals either early or late in infection. At least eighteen bloodstream expression sites were predicted in EATRO 1125, with the initial AnTat1.1 VSG located as an expressed copy on telomere 1iiB (EATRO1125 BES1), a silent second copy also being located on chromosome 8 in inverted orientation, as previously described^37^. Analysis of telomeric VSG genes located in the other EATRO1125 BESs provided evidence of their expression early in infection (day 10- 20), with the respective abundance in the population of BES-resident tVSG expression suggesting a preference for activation of some BES-resident tVSGs over others, either by their differential activation in situ or probability of relocation into EATRO1125 BES1. Strikingly, when clones were analysed from day 17 and day 60-70, the expressed VSG was located in EATRO1125 BES1 in every case and there were only two examples of recombination of VSGs into other expression sites, these not being linked to detectable VSG expression. The VSG present in EATRO1125 BES1 in our clones were also seen to be expressed at equivalent timepoints in vivo across multiple animals. EATRO1125 BES1 is therefore the dominant VSG transcription locus both early and late in infection and other BES-resident VSGs appear to be activated in situ only in a brief window after elimination of the infecting AnTat1.1 VSG or are only then exploited as VSG donors for EATRO1125 BES1. If non-dominant VSG expression sites are transiently activated early in infection this may assist trypanosomes as they establish in a new host to overcome herd immunity ^49^ or to select expression sites optimised for different hosts. Alternatively, in the unusual context of experimental infections with a single infecting antigen type it may be necessary for trypanosomes to activate other expression sites upon immune recognition of the infecting VSG until a new VSG can be successfully recombined into the preferred site. Regardless, the continued use of the dominant site over more than 60 days of infection in cattle highlights that immunity to co-transcribed expression site associated genes is not a selective feature of BES usage. Moreover, the dominant use of EATRO1125 BES1 in both mouse and cow infections supports evidence that there is not a stringent dependence between the dominant ES and host species ^50,51^, contrasting with some previous explanations of expression site diversity and usage^52^ and despite metabolic and gene expression distinctions between parasites in mice and cows ^53^. Interestingly, where mappable, the same telomeric VSGs we observed in non-dominant BESs were at a consistent location in an independently derived EATRO1125 genome ^54^, emphasising that only the dominant BES frequently exchanges VSG. Correspondingly, our clones derived from cows showed little or no evidence of dynamic VSG changes in other expression sites. Hence even if culture adaptation or cloning selected for use of the EATRO 1125 BES, other expression sites appear very stable with little evidence of VSG gene replacement even after more than 60 days in vivo.

Emphasising the importance of mosaic VSG formation throughout chronic trypanosome infections, only 8.5% of the VSG gene archive of EATRO 1125 parasites encoded full length matched transcript sequences detected during the infections. Importantly, our identification criteria required 99% identity over 99% of the intact sequence length, a stringency necessary to unambiguously distinguish the expressed sequence from related VSG sequences in the genome. Of the remaining 91.5% of expressed transcripts, most could be aligned over most of their length using multiple identifiable mosaic donor sequences of at least 50nt in length and with a 3’ end donor bias. This complex diversification of the VSG gene repertoire using identifiable putative donors is compatible with modelling studies of the evolution of the VSG archive. Bayesian approaches have inferred that very short gene conversion tracts of only 14- 25nt (mean 18nt) drive clustered diversification, together with more frequent but distributed point mutations ^55^. Supporting this, experimental analysis of mosaic formation after an induced break in a targeted VSG coding region has identified short homology driven mosaic events^10,12^, consistent with the short (50-100 nt) donor lengths most common in our analysis. Some donor regions could not be identified although microhomology based conversion over short regions, analogous to that observed in the diversification of MHC archive ^56^, renders the unambiguous detection of micromosaic donors less than 50bp in length significantly challenging.

Analysis of VSG expression in cattle has allowed us to capture a much larger number of VSG mosaics than previously described and confirms descriptions of the complex patchwork of donor sequences in any given transcript^18,49^. In addition, the cow data extend our understanding of this process. First, unlike in previous murine studies^18^, we show that VSG mosaic complexity, as measured by the number of donors, increases over the course of an infection, as has been described during AV in Anaplasma infections^57^. Second, we see a notable bias in the location of recombination junctions at the 3’ end of mosaic VSGs, as also noted by PCR-based detection of VSG diversification in mice. One explanation for this bias may lie in the observation that only a small minority (∼15%) of archive VSG genes contain a functional C-terminal domain, which is needed for addition of a glycosyl phosphatidyl inositol anchor to secure the VSG the plasma membrane. Thus, the 3’ gene recombination bias may reflect the necessity to generate C-terminally functional VSGs. However, the VSG 3’ recombination bias may also be genetically programmed, since the location of increased donor usage mirrors recent mapping of a pronounced DNA break within the actively transcribed VSG^10^.

To explore the contribution of known recombination mechanisms to VSG diversification in vivo, VSG expression was examined in null mutants for both BRCA2 and RAD51, whose loss significantly impairs homologous recombination and VSG switching^8,9^. This analysis provided several insights into the operation of AV in long-term *T. brucei* infections. First, we found that after a major first wave of parasitaemia, infections could be sustained by the mutants for over 30 days in mice albeit at reduced parasitaemia potentially linked to the enhanced genome instability generated when RAD51/BRCA2 function is perturbed^40^. Second, gene activation was not restricted to those VSG genes resident in expression sites, demonstrating that VSG gene recombination across the genome is possible in the absence of RAD51-directed homologous recombination, as previously reported ^8,58^. Third, the loss of either RAD51 or BRCA2 did not fully eliminate VSG expression diversity observed both early and late in infection, but the scale of diversity was considerably reduced, and mosaic donor fragment lengths were similar to early infection for wild type parasites. Finally, the VSGs expressed in the two mutants, even at later time points, were hierarchically clustered with those VSGs expressed early in both mice and cow infections by wild type *T. brucei*. In combination, these findings suggest that the absence of homology-directed, RAD51-catalysed, recombination significantly restricts VSG diversity, but there must also be a further contribution from a RAD51-independent mechanism, which remains wholly uncharacterised. Consistent with the predictions from laboratory studies that artificially introduced cleavage within the expressed VSG ^10,12,59,60^ ^61^, this alternative mechanism must, like RAD51-dependent HR, be capable of whole-genome scanning to assemble functional VSGs from related VSG gene donors. Notably, observed subtelomeric deletions in the genome of the recombination mutants may also restrict the available VSG gene archive by the deletion of donor VSGs.

Our results generate a coherent description of the infection dynamics of *T. brucei* in bovine hosts that is consistent with in vitro and rodent models for trypanosome antigen variation but considerably extend our current understanding. This involves use of a dominant VSG expression site both early and late in infection, with only a potentially transient contribution from VSGs in other expression sites in the earliest stages of infection. Indeed, in all analysed clones from different hosts and stages of infection, new antigen expression is highly polarised to a single dominant expression site, which remains active throughout weeks of chronic infection. The diversity of VSG expression is also extreme and unpredictable and yet follows a hierarchical temporal expression of related VSGs, with infections in rodents and bovines following a similar trajectory. This hierarchy is overwhelmingly dominated by macro and micro sequence identity between VSG genes and our analysis of HR mutants reveals that all forms of diversifying VSG recombination during an infection are not the product of one machinery, but at least two: a dominant homologous recombination pathway involving RAD51 and BRCA2, and a further unspecified recombination mechanism. This diversification significantly extends the antigen repertoire available to the trypanosome, supporting the maintenance of chronic infections with sufficient diversity to outcompete the immune system.

## Methods

### Animals

Five Holstein Friesian calves Male (aged 4–6 months) maintained at the Large Animal Research and Imaging Facility of the Roslin Institute, under vector proof containment (BSL2) conditions. To initiate calf infections, blood containing *T. b. brucei* EATRO1125 AnTat 1.1 90:13 was freshly harvested from two infected donor mice, pooled and blood containing 1 × 10^6^ parasites in 1 ml used to infect each calf i.v. Parasitaemias were monitored by microscopy after buffy coat preparation, using a conversion between buffy coat and haemocytometer counts (parasites/ml = 29,478 × parasites/field of view [FOV]) where parasite numbers were at the limit of detection, as described previously^53^. Infections in mice used C57BL/6J male mice at least 12 weeks old, with an inoculum of 1000 *T. brucei* EATRO1125 AnTat 1.1 90:13 parasites. Parasitaemias were determined by the rapid matching method of Herbert and Lumsden^62^. Mouse and cattle experiments were performed using procedures approved under the UK Home Office Animal (Scientific Procedures) Act (1986) license numbers PP2251183 and PP4458006 respectively, each being approved after review at the University of Edinburgh ethical review committee.

### Parasites and parasite culture and cloning

To establish ex vivo cultures from mice, 200µl of whole blood from the terminal bleed was split evenly into 4 wells of a 6 well plate, each containing 5ml HMI9 medium. These cultures were left for at least 96 hours so any remaining parasites committed to cell cycle arrest were removed from the population, ensuring outgrowth of the remaining parasites. The 4 “starter” cultures from each mouse (“pooled populations”) were then used to generate clones: cultures were serially diluted to allow selection of clones where fewer than 1 parasite/well was expected in a 96-well plate. Clones were grown to 2 x 10^8^ cells total, not exceeding 1 x 10^6^/ml density. To pellet the cells, we centrifuged the cultures at 700g, washed twice in PBS, and froze pellets immediately at -80°C for subsequent DNA extraction. Parasites from cattle infections were established ex vivo by diluting whole blood 1/50 in HMI9. Once the cells were doubling at least twice in 24 hours, cells were cloned by dilution in the same way described above. For parasites derived from late cow infections this adaptation process took much longer than those isolated from mice or early cow infections.

### RAD51 and BRCA2 KO

The creation of the mutant lines is detailed in Supplementary Figure 12.

### VSGseq2 and PAD1 qRT PCR sample preparation and sequencing

Throughout WT *T. brucei* mouse infections 20µl volumes of blood were taken in accordance with UK Home office guidelines, and stored at -80°C in 20µl of RNAlater buffer (Thermo Fisher). RNA was subsequently made from the thawed samples using a Kingfisher flex 711 automated extraction robot using MagMax-96 Blood RNA isolation kits (AM1837) with modifications outlined in ^63^. For RNA from parasites from terminal bleeds in mice, the parasites were purified from the blood using DE52 cellulose prior to the pellets beings stored and processed as above. For VSGseq2 we used only the terminal bleeds from WT and mutant mouse infections resulting in one sample per mouse. For cows RNA from longitudinal 5-10ml blood samples was used. cDNA for all samples was generated from RNA using Superscript III or IV one step synthesis kits (Thermo scientific) using oligo dT primers for PAD1 qRT PCR, or VSG specific All VSG 14mer primer outlined in the VSGseq protocol ^14^ for VSGseq2. qRT PCR for PAD1 for WT infections in mice and cows was performed as previously described ^30^ using ZFP3 as a normalisation control. To calculate relative levels of PAD1 in each sample the ΔΔct method was used, where all samples were compared to an average of the d3 values from WT mouse infections, where previous work has shown the population to be composed entirely of fully replicative slender form parasites ^64^. Values are reported as fold changes vs d3 slender using the formula **2^-ΔΔCt^.**

cDNA generated from terminal mouse and longitudinal cow samples was processed according to the VSGseq pipeline ^14^ using primers specific for VSG, after which they were cleaned using AMpure XP beads and sequenced. To compare the efficacy of different sequencing technologies terminal samples from wild type mouse infections were sequenced using both illumina (Hiseq 4000) and PacBIO (Sequel). Using the original VSGseq pipeline^14^, the results suggested equivalent performance such that further VSGseq2 analysis used only the illumina platform (for details of all NGS sequencing for our samples see Supplementary data 1). After sequencing, subsequent analyses were performed according to the VSGseq2 pipeline using default settings ^65^

### DNA extraction and genomic analysis

Frozen cell pellets were thawed and DNA extraction proceeded immediately, using the Monarch HMW DNA Extraction Kit for Blood and Tissues (New England Biolabs) to extract gDNA according to the manufacturers protocol. DNA was resuspended in 100µl of Elution Buffer, and stored at -20°C before being sequenced.

For the starting reference genome, verkko (v2.2.1) ^33^ was used to assemble the genome using HIFI, Nanopore and HI-C data. Quality control was performed with QUAST (v5.3.0) ^66^and BUSCO (v5.7.0_cv1), Genomic data from mouse and cow samples throughout the infection was also assembled; as only HIFI data was available for these samples, flye (v 2.9.6-b1802) was used for assembly.

### Mosaic analysis

To identify potential mosaic VSGs, custom python scripts were utilised to match VSG transcript sequences with genomic ones. First, a genomic kmer (k=50) database was constructed - this comprised four sources of sequence: two genome assemblies from the start of the in vivo experiments (assembled using verkko and flye), the EATRO 1125 reference genome from TriTrypDB (tritrypdb.org/, genome build 68)^54^, and the EATRO 1125 VSG collection available at https://tryps.rockefeller.edu/Sequences.html (CDS with flanking sequences, min. 150 amino acid length). This kmer index was queried using VSG transcript sequences, in overlapping 50 bp bins with a 1 bp step. Adjacent 50mer sequence matches were merged, and, following this, donor identification was performed; first, potential donors that were fully encompassed within other, larger potential sequence donors, were removed. If potential sequence donors were identified within 2kb of each other in the VSGome database, they were considered to belong to the same, single, donor. In situations where donors were found to contribute more than one sequence fragment, an allowance of up to 3bp in gap length was made to consider fragments as belonging to a single fragment (containing a likely polymorphism of up to 3bp).

Transcripts analysed downstream were filtered by minimum coverage (80% or 99%, indicated), and a high-fidelity subset of the data (referred to as ‘high confidence’ transcripts) was defined as a VSG transcript having at least 100-fold coverage present in a minimum of two distinct samples. The mosaic VSG analysis was performed using custom python scripts, utilising the following packages: pandas (v.2.0.3), numpy (1.24.4), scipy (1.10.1), matplotlib (3.7.5), seaborn (0.13.2).

## Data availability

Sequencing data associated with this study has been deposited under the BioProject PRJNA1278329. VSG mosaic data for all transcripts is available at doi.org/10.5281/zenodo.21294599.

## Code

Code has been deposited here: https://github.com/goldrieve/COBALT

## Author contributions

Experiments: SL, JM, CL, CV, EP, LJM

Data analysis: SL, GO, AJ, MK, KRM, CD, MM, RM, MM

Experiment conceptualisation: SL, JM, KRM, LJM, RM, AJ Manuscript preparation/Editing: KRM, SL, JM, MM, AJ, RM, LJM,

## Acknowledgements

We would like to acknowledge Kirsty McWilliam for critical reading of the manuscript and the Roslin Institute Large Animal Research and Imaging Facility staff, in particular James Nixon, Peter Tennant, Chris Proudfoot and Adrian Ritchie, for their invaluable help in running the cattle experiments, and Stefano Guido for veterinary input and guidance.

## Supplementary Figures

**Figure S1**

Individual parasitaemias and PAD1 expression of the late mouse group (L1- L8), and individual parasitaemias and PAD1 expression of cattle infections (animals 900, 908, 914, 924, 929). The grey zone indicates where parasites were at or below the limit of detection by microscopy (∼10^4^/ml). PAD1 expression was normalised to a 100% stumpy population.

**Figure S2**

(a) Hierarchical ordering of VSG relatedness in combined mouse and cow samples throughout infections ordered by time. Relatedness is indicated on the left hand dendogram

(a) Dominant VSGs shared between mouse (WT-early; WT_late) or cow infections (dpi10, 20, 30, 40, 50, 60, 70) at different time points. Each time point represents the combined VSG clusters within all animals at that time point or time period.

(a) Dominant VSGs shared between individual cow infections

**Figure S3**

Plot of VSGs of different lengths against their timing of peak expression

**Figure S4**

Packed cell volume (haematocrit) of each individual cow throughout the time of infections.

**Figure S5**

VSG expression profiles for all individual VSG clusters (labelled in the grey box above each profile) that are expressed in 3 or more animals over time. Y axis=TPM; X axis = days post infection. Each animal is colour coded as indicated at the top of the Figure. Asterisks indicate tVSG present at EATRO 1125 BES1 in isolated clones.

**Figure S6a-e**

Analysis of individual VSG transcripts within individual animals where VSG clusters reappear later in infection, providing evidence for direct re-expression of some VSGs within an animal, or the assembly of related VSGs by mosaic formation.

**Figure S7**

(a) Genome assembly statistics for the *T. brucei* EATRO 1125 AnTat1.1 genome

(b) Assembly mapping of each haplotype in the EATRO 1125 genome.

**Figure S8**

(a) RNAseq analysis of expression of VSGs in clones derived at the end of the experiment (EOX; 66, 70 and 68 days post infection, dpi) from different calves (924; 908 and 914), compared with the expression of VSGs from the cultured pooled parasites from which the clones were derived (Pool).

(b) Map of EATRO 1125 BES1 with resident VSGs in distinct clones from each animal, isolated either early in infection (d11 or d17, corresponding to peak 2 of infection) or at the end of experiment (d66-70).

**Figure S9**

a-c Coverage of VSG transcripts by potential donor sequences within the EATRO 1125 genome. For the high confidence transcript set (‘gold standard’) and all detected VSG transcripts, sequences were searched for potential donor sequences within the EATRO 1125 verkko assembly from this study, in addition to the EATRO 1125 assembly of the Roditi lab and the VSGome dataset for EATRO 1125 from the Cross lab. This aimed to identify any intact VSGs missing from our verkko assembly and to identify the possible sequences available in all available datasets and so identify potential mosaic donors. For over 80% of transcripts donor coverage could be identified at >99% identity from the search set.

**Figure S10**

a. Mosaic formation in the high confidence and all VSG transcript data set for transcripts where 80% of the sequence is covered by identifiable donors. The histogram shows the number of donors per transcript.

b. Representation of the number of fragments contributed by each donor that cover at least 80% of the VSG transcript at different times of infection in cattle for the entire VSG transcript dataset.

c. Donor alignment statistics for transcripts where donors cover >80% or >99% of the transcript length.

**Figure S11**

Length distribution of donor fragments that contribute to mosaic formation of all transcripts at >80% coverage. Alignment statistics for donor contributions to the

mosaic assembly are tabulated below the histogram with either 80% or 99% coverage.

**Figure S12**

Generation and validation of RAD51 and BRCA2 null mutants in pleomorphic T. brucei

a) Map of the integration site for the deletion of RAD51. Alleles were replaced with either Blasticidin or puromycin, with flox excision removing the associated drug resistance genes. PCRs Amplicons primed outside of the coding region (JM024, JM025) or within the coding region (JM 032, JM033) confirmed the correct integration of the selection cassettes and appropriate validation of null mutant clones. Two mutants (1-25; 2-1) were derived

b) Map of the integration site for the deletion of BRCA2. Alleles were replaced with either Blasticidin or puromycin, with flox excision removing the associated drug resistance genes. PCRs Amplicons primed outside of the coding region (JM026, JM027) or within the coding region (JM 034, JM035) confirmed the correct integration of the selection cassettes and appropriate validation of the resulting null mutant clone (B6N1).

c) Confirmation of the absence of RAD51 (blue) or BRCA2 (green) transcripts in the generated null mutants for each gene. Data represent Mean of 2-4 experiments; 3 technical replicates per experiment.

d) In vitro growth of RAD51 and BRCA2 null mutants in comparison to the parental line. Data represents the mean of 2-3 experiments; each with 3-4 replicates.

**Figure S13**

*Top* Analysis of the genomic origin of the top 10 mosaic VSGs expressed by RAD51 mutant parasites. The potential origin of each donor fragment is indicated in the key below.

*Bottom* Representation of the same mosaic VSGs, with donor fragments located at a putative expression site highlighted in red.

**Figure S14**

a. Genomic sequencing analyses for RAD51 (“R1pxcy_fly.fasta’ “R2p*x*c*y*_fly.fasta) and BRCA2 (“B1p*x*c*y*_fly.fasta”) mutants or parental cells derived early or late in infection (“TbMSEp*x*p*y*p*z*E_fly.fasta”; “TbMSLp*x*p*y*p*z*L_fly.fasta”). The overall sequencing quality for all samples was similar.

b. Alignment of the parental, BRCA2 and RAD51 clonal genomes against the verkko assembly of the EATRO1125 starting genome. Although some regions aligned poorly across all genomes, for the RAD51 and BRCA genomes subtelomeric regions are missing for many chromosomes (black bounding box).

## Supplementary data

**Supplementary data 1**

Sequencing technologies and methodologies for the analysis of all nucleic acid sequences derived in this study

**Supplementary data 2**

‘Champion’ VSG sequences representative of each VSG cluster derived by VSGSeq2. See Oldrieve et al, 2025 ^28^ for a description of the derivation of champion VSGs. Essentially, in vsgseq2, cd-hit-est picks a single VSG from a cluster of related VSGs to represent all VSGs within that cluster. The champion VSG has highest mean expression across all samples.

**Supplementary data 3**

VSG cluster transcript abundance (tpm) in each sample from all mice (early and late samples) and all cows (on each sampling day).

