## Supplementary Figure 1-14 for "Restricted expression site use and extreme genome diversification drives trypanosome antigenic variation in chronic bovine infections"

### Supplementary figures

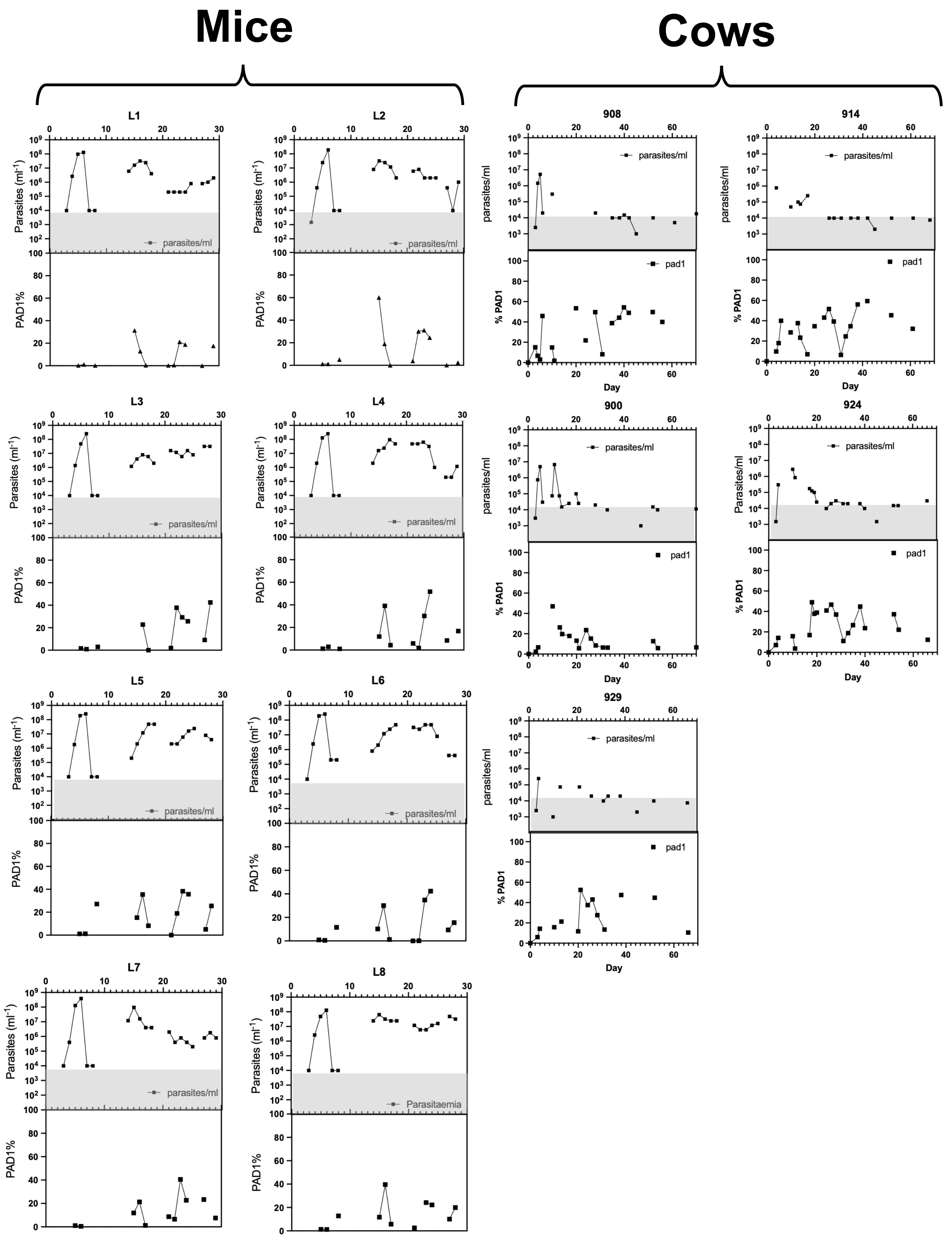
Mice
Cows
L1
L2
L3
L4
L5
L6
L7
L8
908
914
900
924
929
Parasites (ml-1)
PAD1%
parasites/ml
Parasitaemia
pad1
% PAD1
Day

Figure S1

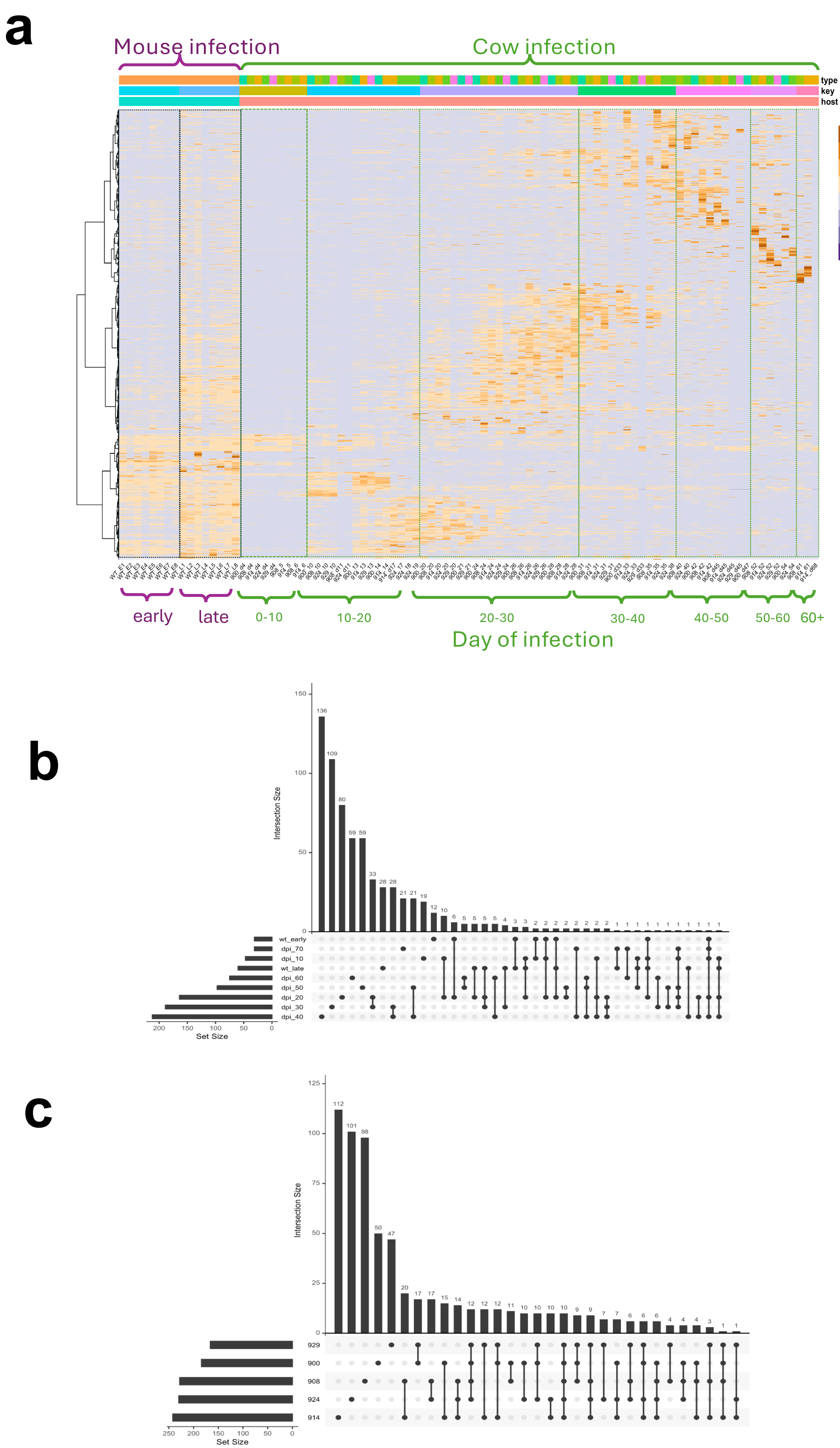
a
Mouse infection
Cow infection
type
key
host
Animal
900
908
914
924
929
WT
Time (days)
0_10
10_20
20_30
30_40
40_50
50_60
60_70
Early
Late
Host
cow
mouse
5
0
-5
early
late
0-10
10-20
20-30
30-40
40-50
50-60
60+
Day of infection
b
Intersection Size
150
100
50
0
136
109
80
59 59
33
28 28
21 21
19
12
10
6
5 5 5 5
4
3 3
2 2 2 2 2 2 2 2
1 1 1 1 1 1 1 1 1 1 1 1
wt_early
dpi_70
dpi_10
wt_late
dpi_60
dpi_50
dpi_20
dpi_30
dpi_40
200 150 100 50 0
Set Size
c
Intersection Size
125
100
75
50
25
0
112
101
98
50
47
20
17 17
15
14
12 12 12
11
10 10 10 10
9 9
7 7
6 6 6
4 4 4
3
1 1
929
900
908
924
914
250 200 150 100 50 0
Set Size

Figure S2 a-c

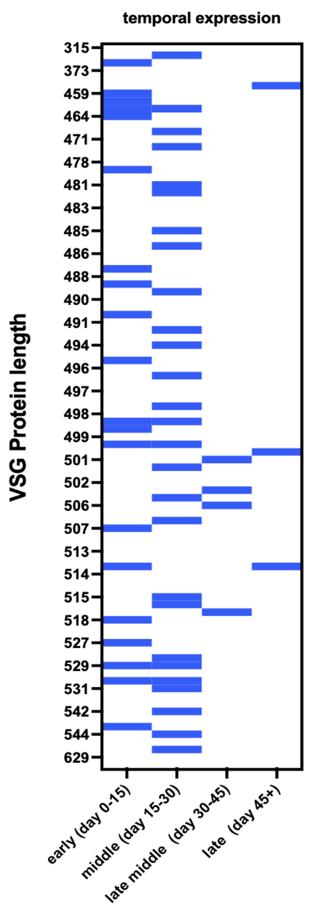
temporal expression
VSG Protein length
315
373
459
464
471
478
481
483
485
486
488
490
491
494
496
497
498
499
501
502
506
507
513
514
515
518
527
529
531
542
544
629
early (day 0-15)
middle (day 15-30)
late middle (day 30-45)
late (day 45+)

### Figure S3

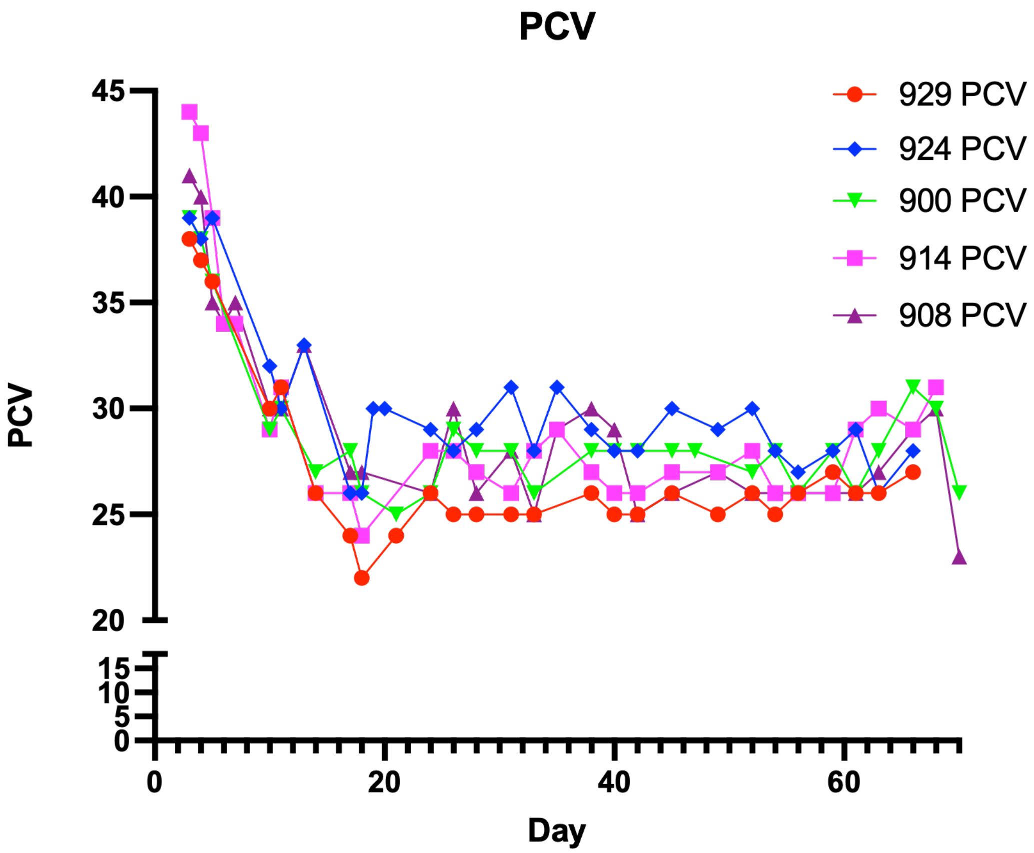
PCV
45
40
35
30
25
20
15
10
5
0
PCV
0
20
40
60
Day
929 PCV
924 PCV
900 PCV
914 PCV
908 PCV

**Figure S4**

### Shared VSGs between ≥3 animals

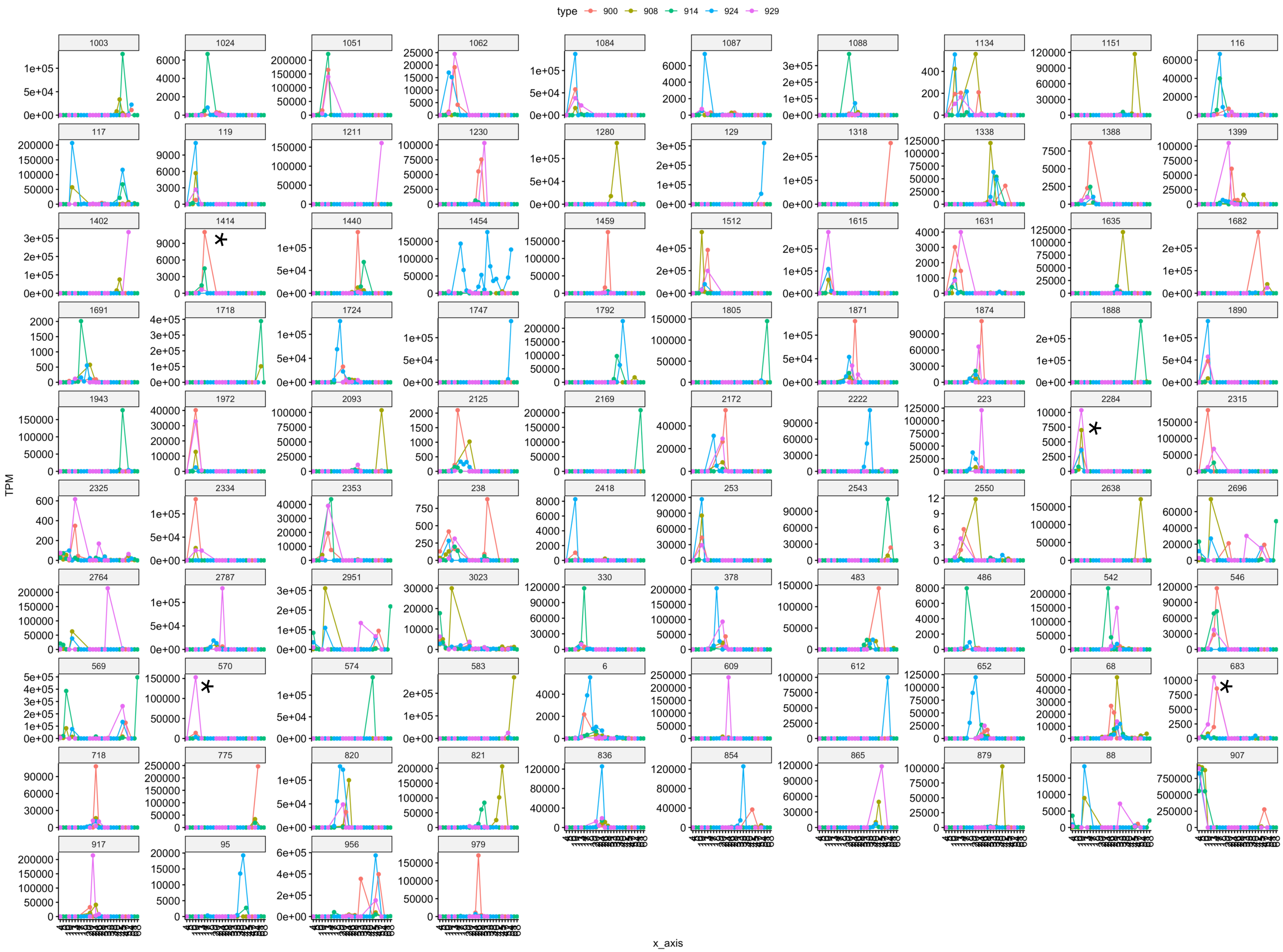

### Figure S5

### AnTat1.1 re-expression: mosaic

#### Cluster 907 cow 900 day 4 expression:

100% match over full length of cluster 907
Detects AnTat1.1. basic copy gene

Descriptions | Graphic Summary | Alignments

**Sequences producing significant alignments** — Download — Select columns — Show 100

select all *3 sequences selected* — Graphics — Distance tree of results — MSA Viewer

| | Description | Scientific Name | Max Score | Total Score | Query Cover | E value | Per. Ident | Acc. Len | Accession |
|---|---|---|---|---|---|---|---|---|---|
| ☑ | 900_d4_trinity.602 | | 2727 | 2727 | 100% | 0.0 | 100.00% | 1512 | Query_7074119 |
| ☑ | 900_d4_trinity.1477 | | 302 | 302 | 13% | 6e-84 | 93.91% | 1428 | Query_7074134 |
| ☑ | 900_d4_trinity.418 | | 168 | 168 | 7% | 3e-43 | 95.24% | 1401 | Query_7074108 |

Sort by description

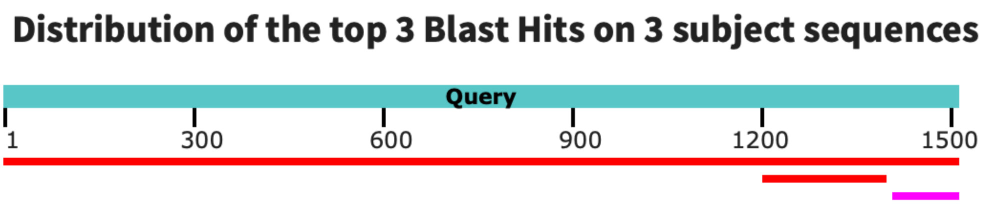

#### Cluster 907 cow 900 day 47 re-expression

73% match over 5' 1096nt; 10% for different transcript at 3' end

Descriptions | Graphic Summary | Alignments

**Sequences producing significant alignments** — Download — Select columns — Show 100

select all *2 sequences selected* — Graphics — Distance tree of results — MSA Viewer

| | Description | Scientific Name | Max Score | Total Score | Query Cover | E value | Per. Ident | Acc. Len | Accession |
|---|---|---|---|---|---|---|---|---|---|
| ☑ | 900_d47_trinity.134 | | 1975 | 1975 | 73% | 0.0 | 99.91% | 1098 | Query_6450368 |
| ☑ | 900_d47_trinity.135 | | 257 | 257 | 10% | 1e-71 | 98.64% | 1509 | Query_6450369 |

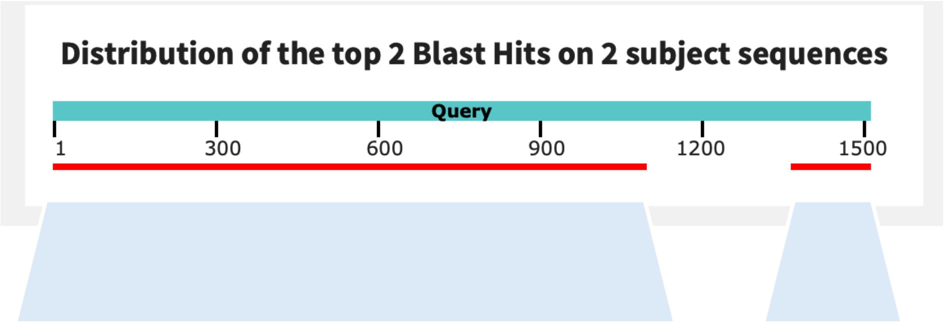

900_d47_trinity.134    900_d47_trinity.135

**Figure S6a**

### Cluster 117 re-expression: non mosaic

Cluster 117, cow 924 day 11

100% match over full length

Descriptions | Graphic Summary | Alignments

**Sequences producing significant alignments** — Download — Select columns — Show 100

select all *16 sequences selected* — Graphics — Distance tree of results — MSA Viewer

| | Description | Scientific Name | Max Score | Total Score | Query Cover | E value | Per. Ident | Acc. Len | Accession |
|---|---|---|---|---|---|---|---|---|---|
| ☑ | 924_d11_trinity.19 | | 2890 | 2890 | 100% | 0.0 | 100.00% | 1602 | Query_5713428 |
| ☑ | 924_d11_trinity.23 | | 181 | 181 | 8% | 9e-48 | 89.63% | 1338 | Query_5713432 |
| ☑ | 924_d11_trinity.22 | | 181 | 181 | 8% | 9e-48 | 89.63% | 1437 | Query_5713431 |
| ☑ | 924_d11_trinity.26 | | 151 | 151 | 7% | 2e-38 | 90.18% | 1536 | Query_5713435 |
| ☑ | 924_d11_trinity.24 | | 87.8 | 117 | 7% | 2e-19 | 78.64% | 1335 | Query_5713433 |
| ☑ | 924_d11_trinity.21 | | 87.8 | 117 | 7% | 2e-19 | 78.64% | 1434 | Query_5713430 |
| ☑ | 924_d11_trinity.4 | | 38.3 | 38.3 | 2% | 2e-04 | 90.00% | 1536 | Query_5713421 |
| ☑ | 924_d11_trinity.3 | | 38.3 | 38.3 | 2% | 2e-04 | 90.00% | 903 | Query_5713420 |

**Distribution of the top 18 Blast Hits on 16 subject sequences**

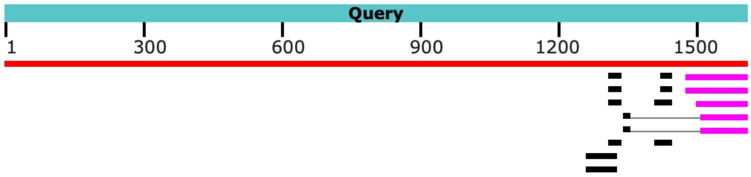

Cluster 117, cow 924 day 45

100% match over full length

Descriptions | Graphic Summary | Alignments

**Sequences producing significant alignments** — Download — Select columns — Show 100

select all *3 sequences selected* — Graphics — Distance tree of results — MSA Viewer

| | Description | Scientific Name | Max Score | Total Score | Query Cover | E value | Per. Ident | Acc. Len | Accession |
|---|---|---|---|---|---|---|---|---|---|
| ☑ | 924_d45_trinity.171 | | 2890 | 2890 | 100% | 0.0 | 100.00% | 1602 | Query_7633371 |
| ☑ | 924_d45_trinity.168 | | 136 | 136 | 7% | 1e-34 | 88.57% | 1509 | Query_7633368 |
| ☑ | 924_d45_trinity.167 | | 122 | 217 | 10% | 2e-30 | 86.27% | 1314 | Query_7633367 |

**Distribution of the top 6 Blast Hits on 3 subject sequences**

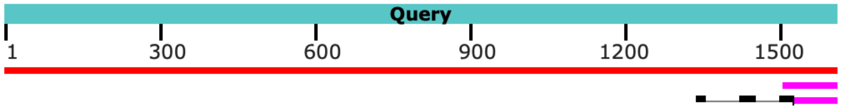

**Figure S6b**

### Cluster 1454: re-expression

#### Cluster 1454, cow 924 day 17;

Descriptions | Graphic Summary | Alignments

**Sequences producing significant alignments** Download Select columns Show 100

select all *96 sequences selected* Graphics Distance tree of results MSA Viewer

| Description | Scientific Name | Max Score | Total Score | Query Cover | E value | Per. Ident | Acc. Len | Accession |
|---|---|---|---|---|---|---|---|---|
| 924_17_trinity.1067 | | 2642 | 2642 | 100% | 0.0 | 99.93% | 1467 | Query_4984000 |
| 924_17_trinity.1069 | | 2636 | 2636 | 100% | 0.0 | 99.80% | 1470 | Query_4984002 |
| 924_17_trinity.1068 | | 2633 | 2633 | 100% | 0.0 | 99.80% | 1470 | Query_4984001 |
| 924_17_trinity.1514 | | 241 | 241 | 10% | 2e-64 | 97.22% | 1536 | Query_4984129 |
| 924_17_trinity.1513 | | 241 | 241 | 10% | 2e-64 | 97.22% | 1536 | Query_4984128 |
| 924_17_trinity.1512 | | 241 | 241 | 10% | 2e-64 | 97.22% | 1536 | Query_4984127 |
| 924_17_trinity.1065 | | 209 | 209 | 10% | 9e-55 | 92.36% | 1542 | Query_4983998 |

**Distribution of the top 100 Blast Hits on 96 subject sequenc**

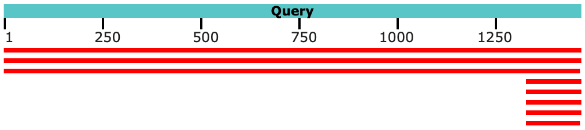

#### Cluster 1454 Cow 924 day 33:

Descriptions | Graphic Summary | Alignments

**Sequences producing significant alignments** Download Select columns Show 100

select all *63 sequences selected* Graphics Distance tree of results MSA Viewer

| Description | Scientific Name | Max Score | Total Score | Query Cover | E value | Per. Ident | Acc. Len | Accession |
|---|---|---|---|---|---|---|---|---|
| 924_33_trinity.2132 | | 2646 | 2646 | 100% | 0.0 | 100.00% | 1467 | Query_5606587 |
| 924_33_trinity.2130 | | 255 | 255 | 10% | 7e-69 | 100.00% | 1542 | Query_5606585 |
| 924_33_trinity.2129 | | 255 | 255 | 10% | 7e-69 | 100.00% | 1542 | Query_5606584 |
| 924_33_trinity.2128 | | 255 | 255 | 10% | 7e-69 | 100.00% | 1530 | Query_5606583 |

**Distribution of the top 63 Blast Hits on 63 subject sequences**

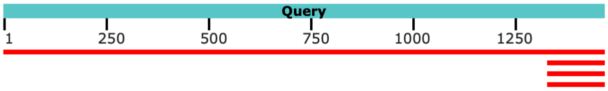

#### Cluster 1454 cow 924 day 54

Descriptions | Graphic Summary | Alignments

**Sequences producing significant alignments** Download Select columns Show 100

select all *42 sequences selected* Graphics Distance tree of results MSA Viewer

| Description | Scientific Name | Max Score | Total Score | Query Cover | E value | Per. Ident | Acc. Len | Accession |
|---|---|---|---|---|---|---|---|---|
| 924_54_trinity.1517 | | 2646 | 2646 | 100% | 0.0 | 100.00% | 1467 | Query_4750903 |
| 924_54_trinity.1133 | | 178 | 178 | 10% | 7e-46 | 87.50% | 1494 | Query_4750890 |
| 924_54_trinity.2263 | | 144 | 144 | 10% | 1e-35 | 83.33% | 1536 | Query_4750952 |

**Distribution of the top 42 Blast Hits on 42 subject sequences**

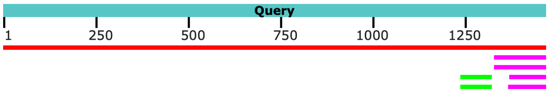

**Figure S6c**

### Cluster 956: reexpression (900); mosaic (914)

#### Cluster 956, cow 900 day 33;

Descriptions | Graphic Summary | Alignments

**Sequences producing significant alignments** Download Select columns Show 100

select all *50 sequences selected* Graphics Distance tree of results MSA Viewer

| Description | Scientific Name | Max Score | Total Score | Query Cover | E value | Per. Ident | Acc. Len | Accession |
|---|---|---|---|---|---|---|---|---|
| 900_d33_trinity.14 | | 2722 | 2722 | 100% | 0.0 | 100.00% | 1509 | Query_8154101 |
| 900_d33_trinity.15 | | 2700 | 2700 | 100% | 0.0 | 99.67% | 1509 | Query_8154102 |
| 900_d33_trinity.1210 | | 210 | 210 | 12% | 2e-55 | 85.56% | 1512 | Query_8154314 |

##### Distribution of the top 50 Blast Hits on 50 subject sequences

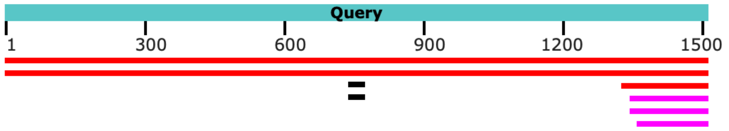

#### Cluster 956 Cow 900 d47 : 100% match

Descriptions | Graphic Summary | Alignments

**Sequences producing significant alignments** Download Select columns Show 100

select all *1 sequences selected* Graphics Distance tree of results MSA Viewer

| Description | Scientific Name | Max Score | Total Score | Query Cover | E value | Per. Ident | Acc. Len | Accession |
|---|---|---|---|---|---|---|---|---|
| 900_d47_trinity.135 | | 2722 | 2722 | 100% | 0.0 | 100.00% | 1509 | Query_446543 |

##### Distribution of the top 1 Blast Hits on 1 subject sequences

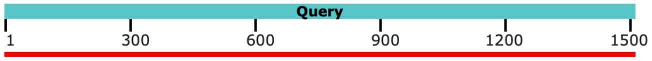

#### Cluster 956 cow 914 day 17: 99.93% match (short stretch of ambiguity)

Descriptions | Graphic Summary | Alignments

**Sequences producing significant alignments** Download Select columns Show 100

select all *100 sequences selected* Graphics Distance tree of results MSA Viewer

| Description | Scientific Name | Max Score | Total Score | Query Cover | E value | Per. Ident | Acc. Len | Accession |
|---|---|---|---|---|---|---|---|---|
| 914_d17_trinity.25 | | 2718 | 2718 | 100% | 0.0 | 99.93% | 1509 | Query_1125952 |
| 914_d17_trinity.24 | | 2718 | 2718 | 100% | 0.0 | 99.93% | 1509 | Query_1125951 |
| 914_d17_trinity.379 | | 248 | 248 | 10% | 1e-66 | 97.28% | 1539 | Query_1126126 |

##### Distribution of the top 107 Blast Hits on 100 subject sequences

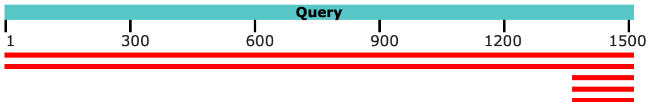

#### Cluster 956 cow 914 day 45:

Descriptions | Graphic Summary | Alignments

**Sequences producing significant alignments** Download Select columns Show 100

select all *93 sequences selected* Graphics Distance tree of results MSA Viewer

| Description | Scientific Name | Max Score | Total Score | Query Cover | E value | Per. Ident | Acc. Len | Accession |
|---|---|---|---|---|---|---|---|---|
| 914_d45_trinity.700 | | 229 | 229 | 11% | 8e-61 | 91.82% | 1533 | Query_3837774 |
| 914_d45_trinity.699 | | 229 | 229 | 11% | 8e-61 | 91.82% | 1533 | Query_3837773 |
| [illegible] | | [illegible] | [illegible] | [illegible] | [illegible] | [illegible] | [illegible] | Query_3837614 |

##### Distribution of the top 103 Blast Hits on 93 subject sequences

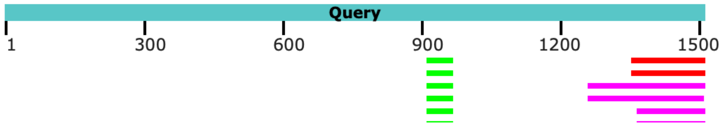

**Figure S6d**

**a**

| | TREU_927 | Hifiasm/purge_dups | Verkko | Verkko_hap 1 | Verkko_hap 2 |
|---|---|---|---|---|---|
| Contigs (N) | 131 | 35 | 54 | 11 | 15 |
| Length | 35,826,294 | 42,706,083 | 83,221,090 | 34,591,865 | 40,198,098 |
| Largest contig | 5,598,354 | 7,731,486 | 7,747,796 | 7,747,796 | 6,701,293 |
| N50 | 3,542,885 | 4,883,410 | 3,457,190 | 3,819,983 | 3,457,190 |
| GC % | 45.49 | 43.67 | 44.21 | 44.87 | 44.25 |
| BUSCO (C) | 99.3 | 100 | 100 | 98.5 | 99.3 |
| BUSCO (S) | 90.8 | 99.2 | 0.8 | 98.5 | 98.5 |
| BUSCO (D) | 8.5 | 0.8 | 99.2 | 0 | 0.8 |
| BUSCO (F) | 0 | 0 | 0 | 0 | 0 |
| BUSCO (M) | 0.7 | 0 | 0 | 1.5 | 0.7 |

**b**

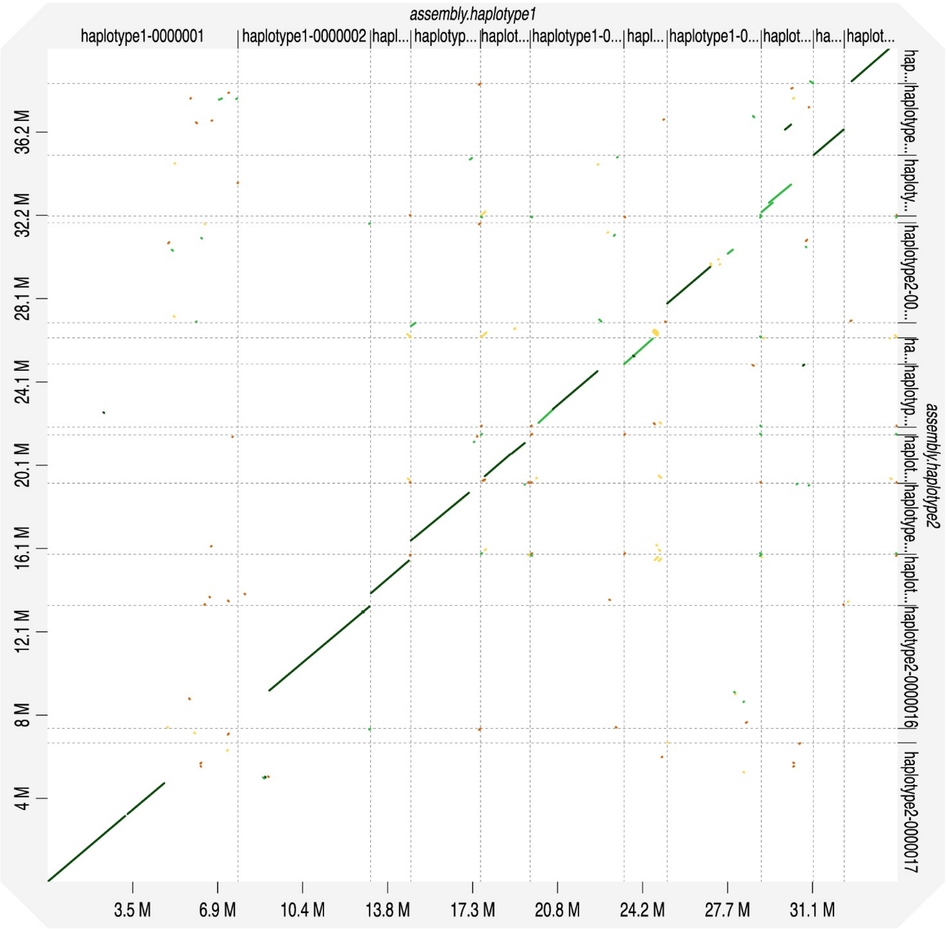

**Figure S7**

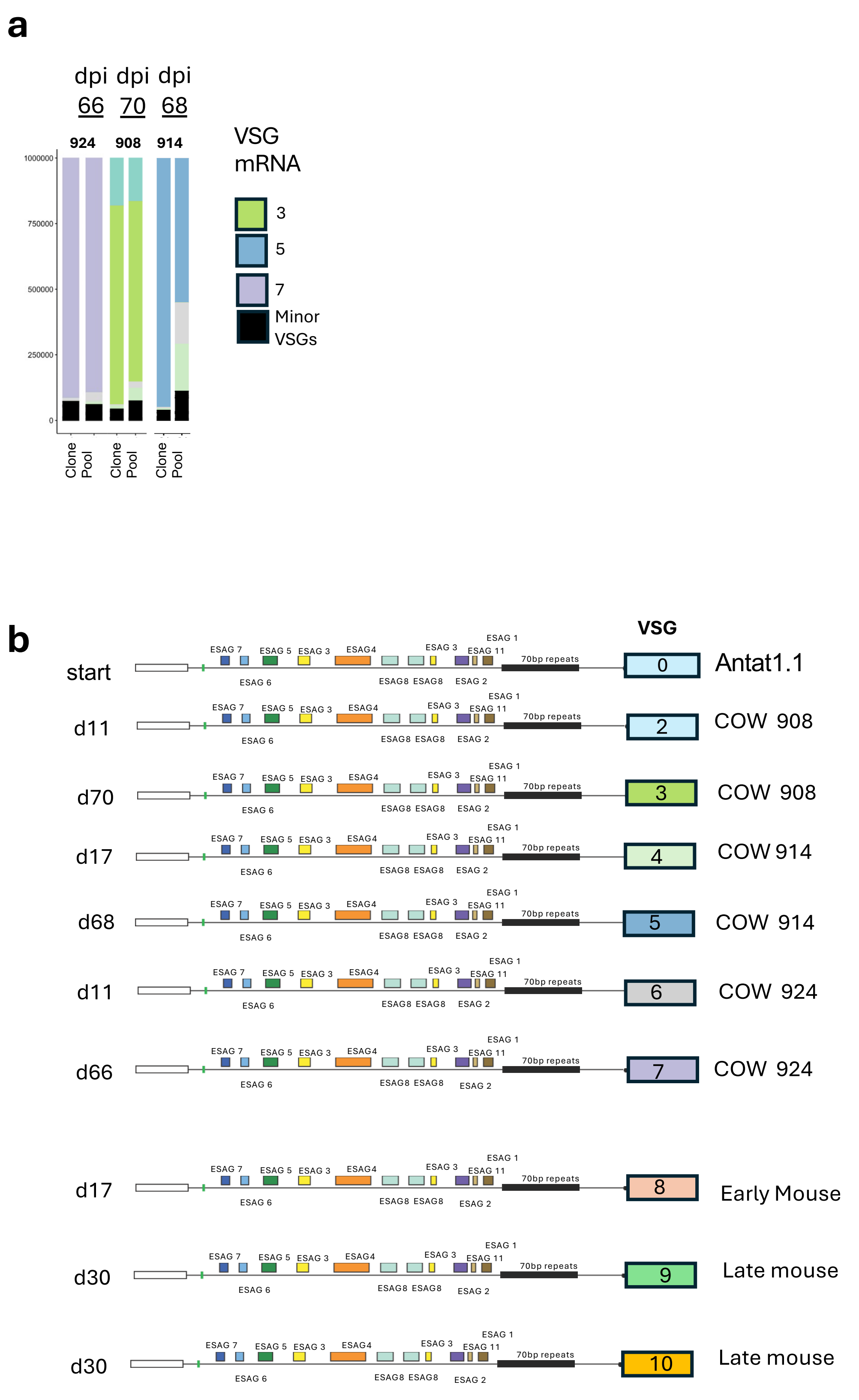
a
dpi dpi dpi
66 70 68
924 908 914
1000000
750000
500000
250000
0
Clone
Pool
Clone
Pool
Clone
Pool
VSG mRNA
3
5
7
Minor VSGs
b
VSG
ESAG 7
ESAG 5
ESAG 3
ESAG4
ESAG 3
ESAG 1
ESAG 11
70bp repeats
ESAG 6
ESAG8 ESAG8
ESAG 2
start
0
Antat1.1
d11
2
COW 908
d70
3
COW 908
d17
4
COW 914
d68
5
COW 914
d11
6
COW 924
d66
7
COW 924
d17
8
Early Mouse
d30
9
Late mouse
d30
10
Late mouse

**Figure S8**

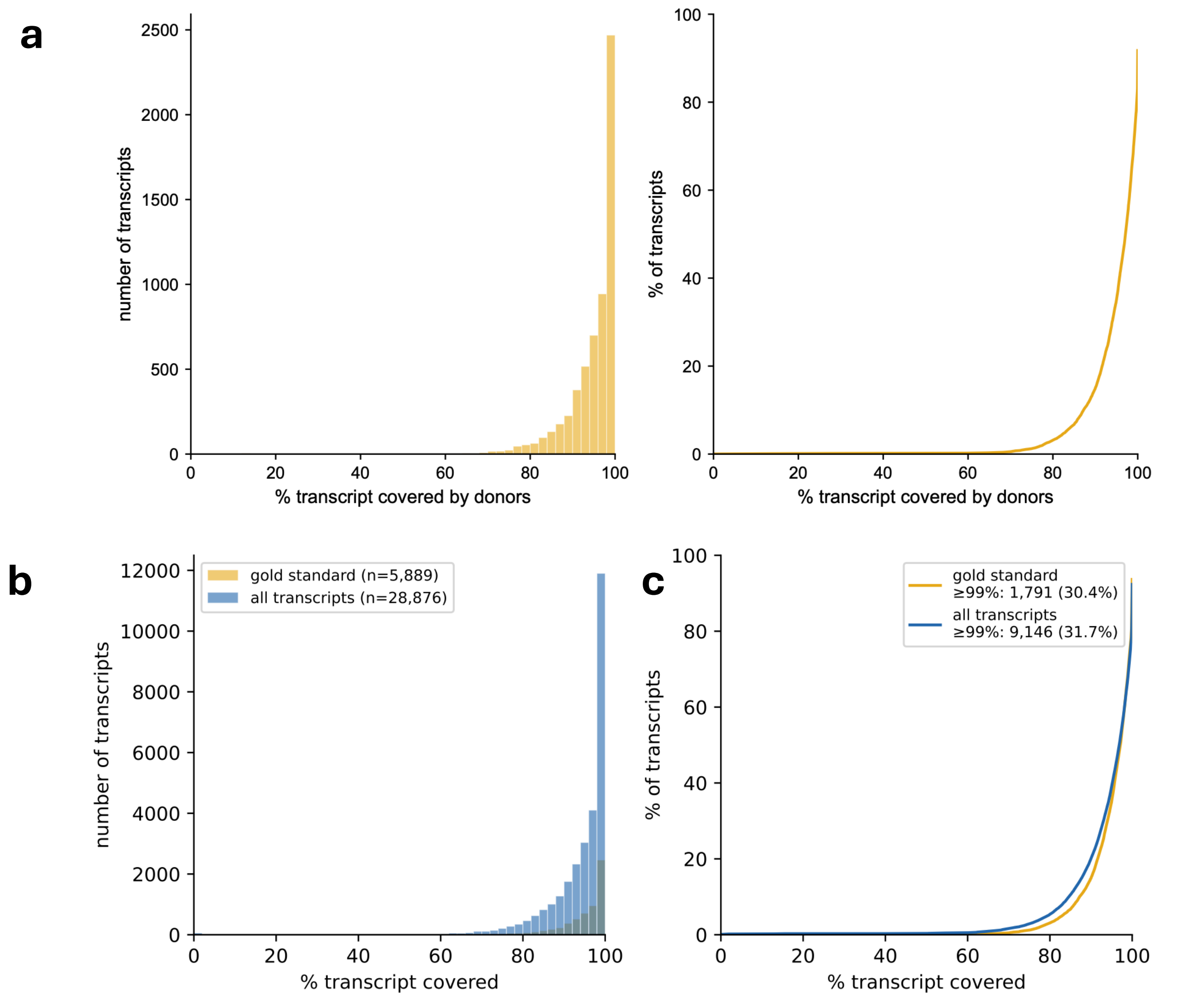

| High confidence (n=5,889, 0%: 1) | |
|---|---|
| count | 5889.00 |
| mean | 95.02 |
| std | 6.60 |
| min | 0.00 |
| 5% | 82.99 |
| 25% | 93.00 |
| 50% | 97.12 |
| 75% | 99.42 |
| 95% | 100.00 |
| max | 100.00 |

| All transcripts (n=28,876, 0%: 56) | |
|---|---|
| count | 28876.00 |
| mean | 94.04 |
| std | 8.66 |
| min | 0.00 |
| 5% | 79.43 |
| 25% | 91.67 |
| 50% | 96.86 |
| 75% | 99.54 |
| 95% | 100.00 |
| max | 100.00 |

**Figure S9**

**a**

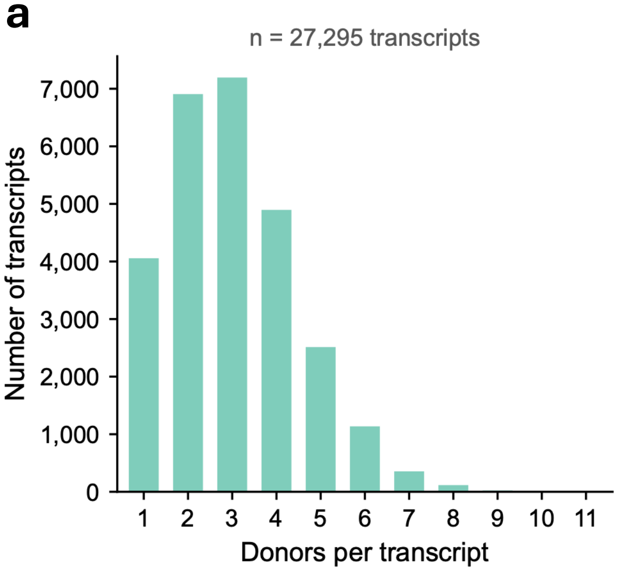

**b**

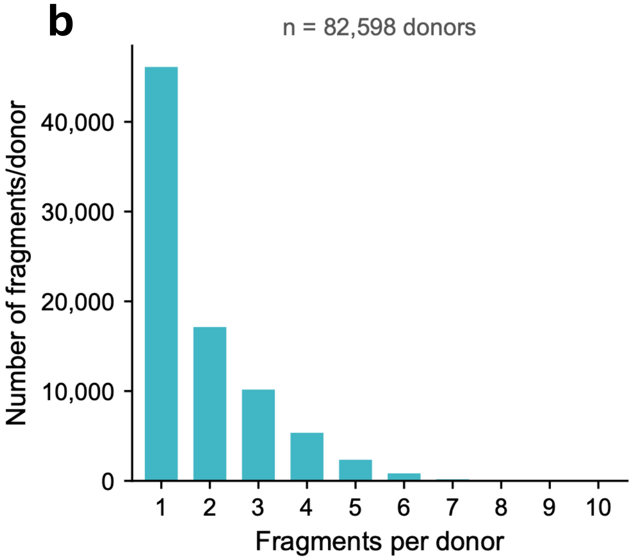

**c**

| | **min. 80% coverage** | **min. 99% coverage** |
|---|---|---|
| | n_donors per transcript | |
| count | 27295 | 9094 |
| mean | 3.03 | 2.42 |
| std | 1.47 | 1.28 |
| min | 1 | 1 |
| 25% | 2 | 1 |
| 50% | 3 | 2 |
| 75% | 4 | 3 |
| 95% | 6 | 5 |
| max | 11 | 10 |
| count | 82598 | 22050 |
| mean | 1.84 | 1.46 |
| std | 1.2 | 0.8 |
| min | 1 | 1 |
| 25% | 1 | 1 |
| 50% | 1 | 1 |
| 75% | 2 | 2 |
| 95% | 4 | 3 |
| max | 10 | 7 |

**Figure S10**

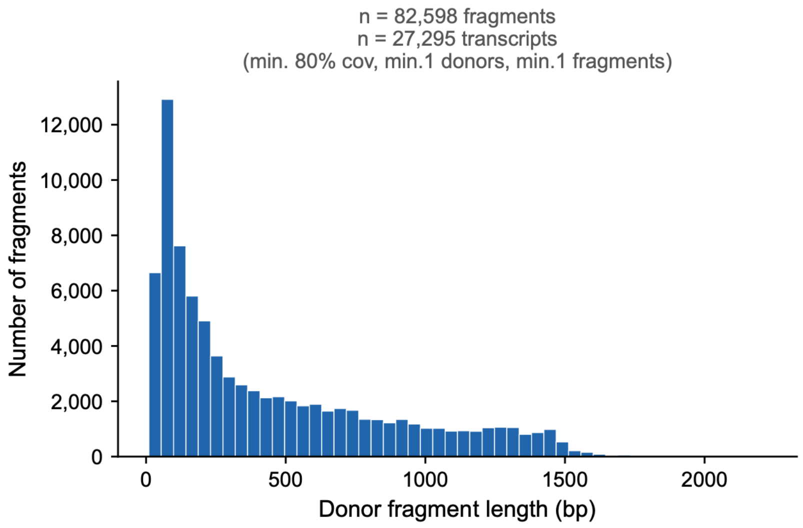

| | min. 80% coverage | min. 99% coverage |
|---|---|---|
| **Fragment length** | | |
| count | 82598 | 22050 |
| mean | 442.3 | 579 |
| std | 414 | 500.5 |
| min | 10 | 10 |
| 5% | 48 | 46 |
| 25% | 104 | 119 |
| 50% | 272 | 408 |
| 75% | 695 | 1013 |
| 95% | 1320 | 1452 |
| max | 2220 | 2220 |

**Figure S11**

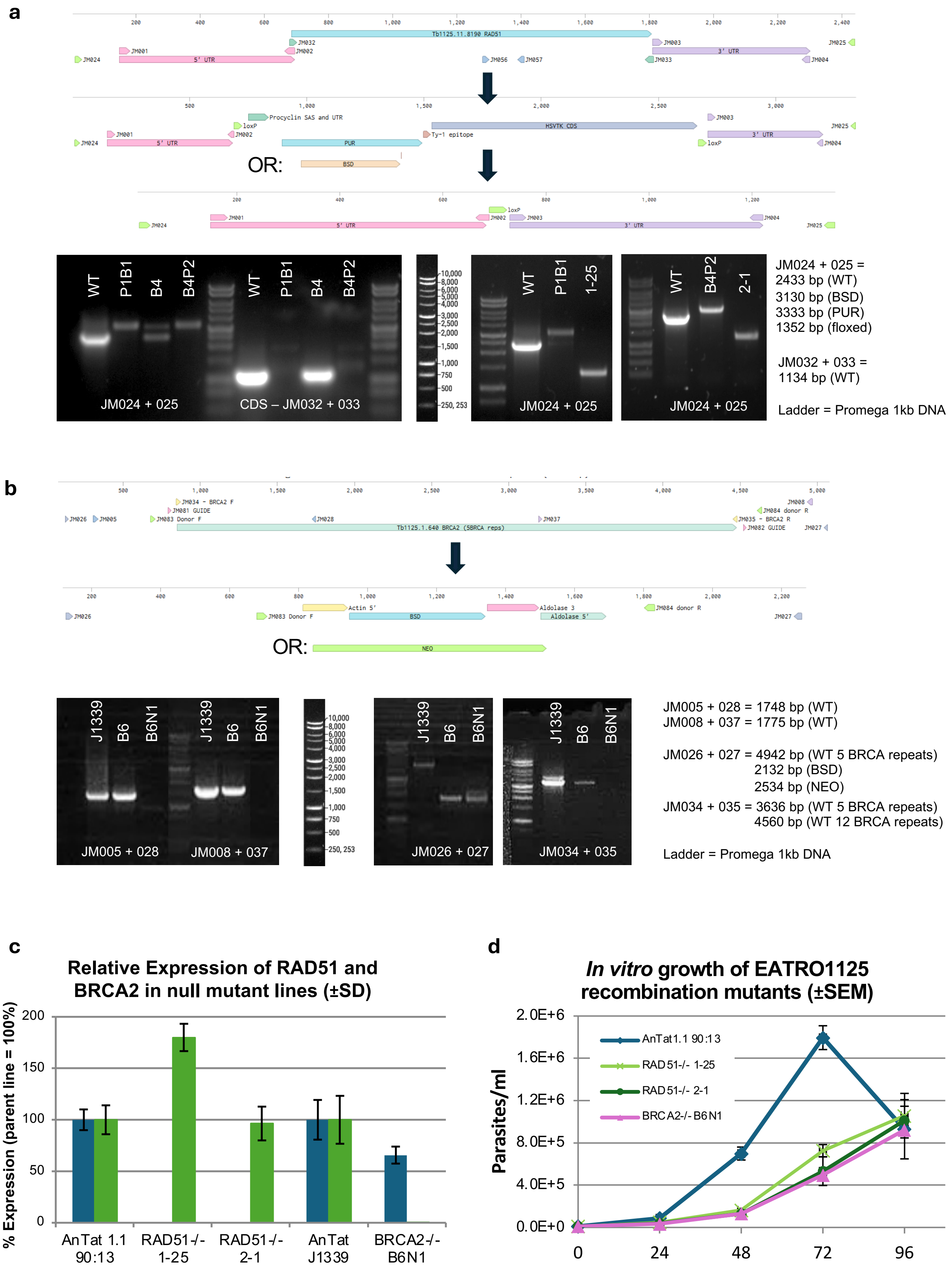
a
OR:
WT
P1B1
B4
B4P2
JM024 + 025
CDS – JM032 + 033
1-25
2-1
10,000
8,000
6,000
5,000
4,000
3,000
2,500
2,000
1,500
1,000
750
500
250, 253
JM024 + 025 =
2433 bp (WT)
3130 bp (BSD)
3333 bp (PUR)
1352 bp (floxed)
JM032 + 033 =
1134 bp (WT)
Ladder = Promega 1kb DNA
b
OR:
J1339
B6
B6N1
JM005 + 028
JM008 + 037
JM026 + 027
JM034 + 035
JM005 + 028 = 1748 bp (WT)
JM008 + 037 = 1775 bp (WT)
JM026 + 027 = 4942 bp (WT 5 BRCA repeats)
2132 bp (BSD)
2534 bp (NEO)
JM034 + 035 = 3636 bp (WT 5 BRCA repeats)
4560 bp (WT 12 BRCA repeats)
Ladder = Promega 1kb DNA
c
Relative Expression of RAD51 and BRCA2 in null mutant lines (±SD)
% Expression (parent line = 100%)
200
150
100
50
0
AnTat 1.1 90:13
RAD51-/- 1-25
RAD51-/- 2-1
AnTat J1339
BRCA2-/- B6N1
d
In vitro growth of EATRO1125 recombination mutants (±SEM)
Parasites/ml
2.0E+6
1.6E+6
1.2E+6
8.0E+5
4.0E+5
0.0E+0
0
24
48
72
96
AnTat1.1 90:13
RAD51-/- 1-25
RAD51-/- 2-1
BRCA2-/- B6N1

### Figure S12

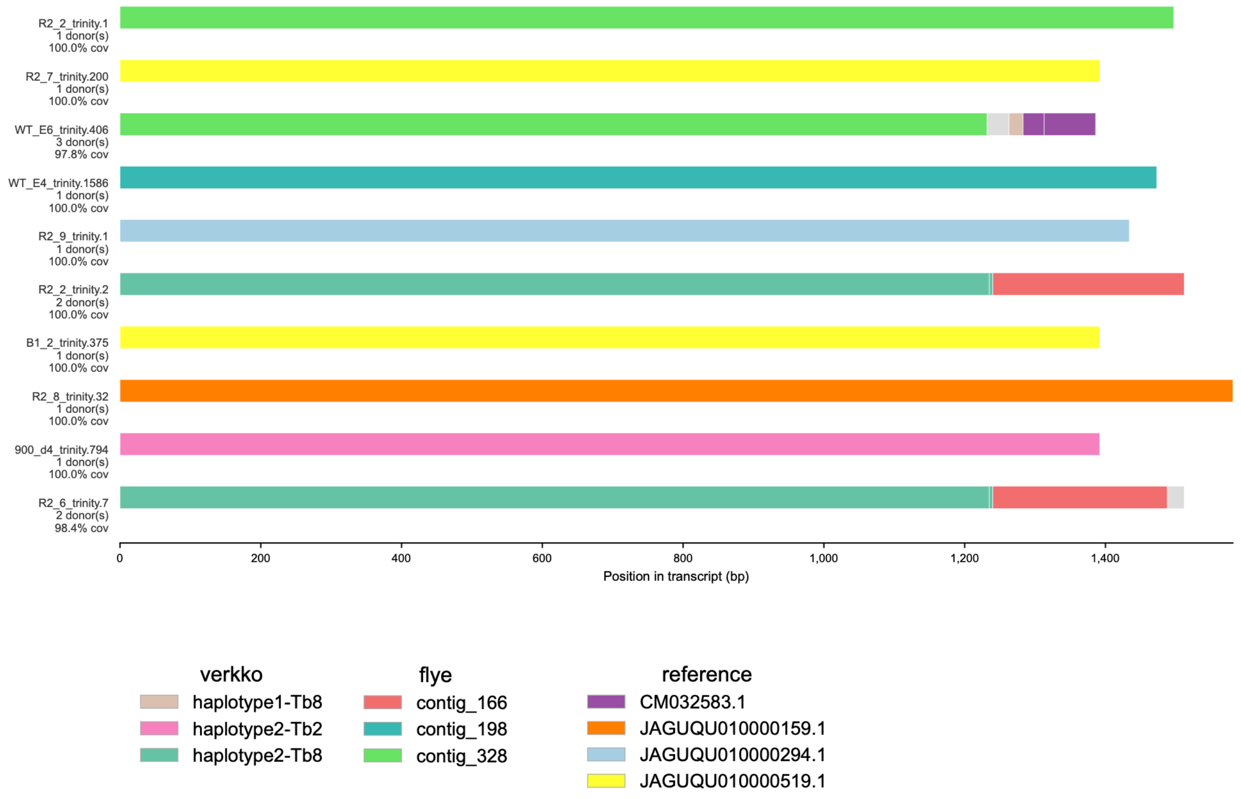
R2_2_trinity.1
1 donor(s)
100.0% cov
R2_7_trinity.200
1 donor(s)
100.0% cov
WT_E6_trinity.406
3 donor(s)
97.8% cov
WT_E4_trinity.1586
1 donor(s)
100.0% cov
R2_9_trinity.1
1 donor(s)
100.0% cov
R2_2_trinity.2
2 donor(s)
100.0% cov
B1_2_trinity.375
1 donor(s)
100.0% cov
R2_8_trinity.32
1 donor(s)
100.0% cov
900_d4_trinity.794
1 donor(s)
100.0% cov
R2_6_trinity.7
2 donor(s)
98.4% cov
0
200
400
600
800
1,000
1,200
1,400
Position in transcript (bp)
verkko
haplotype1-Tb8
haplotype2-Tb2
haplotype2-Tb8
flye
contig_166
contig_198
contig_328
reference
CM032583.1
JAGUQU010000159.1
JAGUQU010000294.1
JAGUQU010000519.1

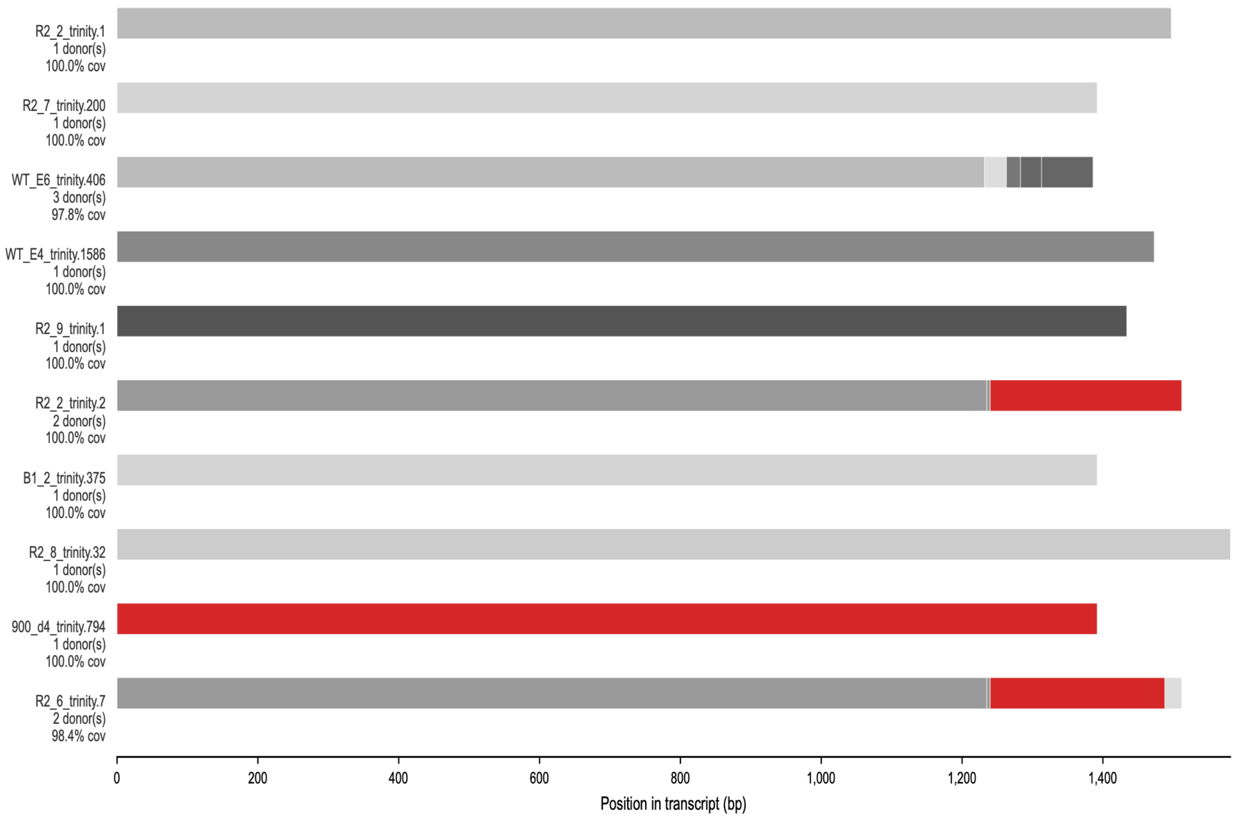
R2_2_trinity.1
1 donor(s)
100.0% cov
R2_7_trinity.200
1 donor(s)
100.0% cov
WT_E6_trinity.406
3 donor(s)
97.8% cov
WT_E4_trinity.1586
1 donor(s)
100.0% cov
R2_9_trinity.1
1 donor(s)
100.0% cov
R2_2_trinity.2
2 donor(s)
100.0% cov
B1_2_trinity.375
1 donor(s)
100.0% cov
R2_8_trinity.32
1 donor(s)
100.0% cov
900_d4_trinity.794
1 donor(s)
100.0% cov
R2_6_trinity.7
2 donor(s)
98.4% cov
0
200
400
600
800
1,000
1,200
1,400
Position in transcript (bp)

**Figure S13**

| Group | Input_file | Dataset | Complete | Single | Duplicated | Fragmented | Missing | n_markers | Internal stop codon percent | Scaffold N50 | Contigs N50 | Percent gaps | Number of scaffolds |
|---|---|---|---|---|---|---|---|---|---|---|---|---|---|
| Starting genomes | AnTat9013start_flye.fasta | trypanosoma_odb12 | 98.1 | 37.7 | 60.4 | 0.3 | 1.6 | 5397 | 0.3 | 726826 | 726826 | 0.00% | 292 |
| | tb_assembly.fasta | trypanosoma_odb12 | 98.1 | 0.2 | 97.9 | 0.2 | 1.6 | 5397 | 0.3 | 3457190 | 3169731 | 0.25% | 54 |
| BRCA2dKO | B1p1c1_flye.fasta | trypanosoma_odb12 | 98 | 37.4 | 60.6 | 0.3 | 1.7 | 5397 | 0.3 | 647059 | 647059 | 0.00% | 302 |
| | B1p5c1_flye.fasta | trypanosoma_odb12 | 97.9 | 37.9 | 60.1 | 0.4 | 1.7 | 5397 | 0.3 | 669416 | 669416 | 0.00% | 331 |
| | B1p5c12_flye.fasta | trypanosoma_odb12 | 98 | 37.6 | 60.4 | 0.3 | 1.7 | 5397 | 0.3 | 618212 | 618212 | 0.00% | 311 |
| RAD51dKO | R1p1c1_flye.fasta | trypanosoma_odb12 | 98.1 | 38.1 | 59.9 | 0.4 | 1.6 | 5397 | 0.3 | 600787 | 600787 | 0.00% | 292 |
| | R1p1c2_flye.fasta | trypanosoma_odb12 | 98 | 36.8 | 61.2 | 0.3 | 1.7 | 5397 | 0.2 | 553850 | 553850 | 0.00% | 309 |
| | R1p4c1_flye.fasta | trypanosoma_odb12 | 98 | 36.5 | 61.5 | 0.3 | 1.7 | 5397 | 0.3 | 601324 | 601324 | 0.00% | 282 |
| | R1p4c2_flye.fasta | trypanosoma_odb12 | 98.1 | 36.8 | 61.3 | 0.3 | 1.6 | 5397 | 0.3 | 539606 | 539606 | 0.00% | 301 |
| | R2p1c1_flye.fasta | trypanosoma_odb12 | 98.1 | 37.9 | 60.2 | 0.3 | 1.6 | 5397 | 0.3 | 607082 | 607082 | 0.00% | 310 |
| | R2p1c2_flye.fasta | trypanosoma_odb12 | 98 | 37.1 | 60.9 | 0.4 | 1.6 | 5397 | 0.3 | 630010 | 630010 | 0.00% | 307 |
| | R2p8c2_flye.fasta | trypanosoma_odb12 | 98 | 37.8 | 60.3 | 0.4 | 1.6 | 5397 | 0.2 | 531289 | 531289 | 0.00% | 313 |
| WT | TbMsE2p3p7p3E_flye.fasta | trypanosoma_odb12 | 98.2 | 34.9 | 63.2 | 0.3 | 1.5 | 5397 | 0.3 | 649547 | 649547 | 0.00% | 290 |
| | TbMsE2p4p8p4E_flye.fasta | trypanosoma_odb12 | 98.1 | 37.6 | 60.5 | 0.3 | 1.7 | 5397 | 0.3 | 566489 | 566489 | 0.00% | 300 |
| | TbMsE2p4p9p7E_flye.fasta | trypanosoma_odb12 | 98.1 | 36.4 | 61.7 | 0.3 | 1.6 | 5397 | 0.3 | 708704 | 708704 | 0.00% | 299 |
| | TbMsL1p16p2L_flye.fasta | trypanosoma_odb12 | 98.1 | 36.2 | 61.8 | 0.3 | 1.6 | 5397 | 0.3 | 652095 | 652095 | 0.00% | 267 |
| | TbMsL1p1p8p2L_flye.fasta | trypanosoma_odb12 | 98.2 | 37.9 | 60.3 | 0.3 | 1.5 | 5397 | 0.3 | 593773 | 593773 | 0.00% | 322 |
| | TbMsL5p2p4p2L_flye.fasta | trypanosoma_odb12 | 98.1 | 34.6 | 63.5 | 0.3 | 1.6 | 5397 | 0.3 | 588022 | 588022 | 0.00% | 302 |
| | TbMsL5p3p6p2L_flye.fasta | trypanosoma_odb12 | 98.1 | 36.1 | 61.9 | 0.3 | 1.6 | 5397 | 0.2 | 684852 | 684852 | 0.00% | 286 |

**Figure S14a**

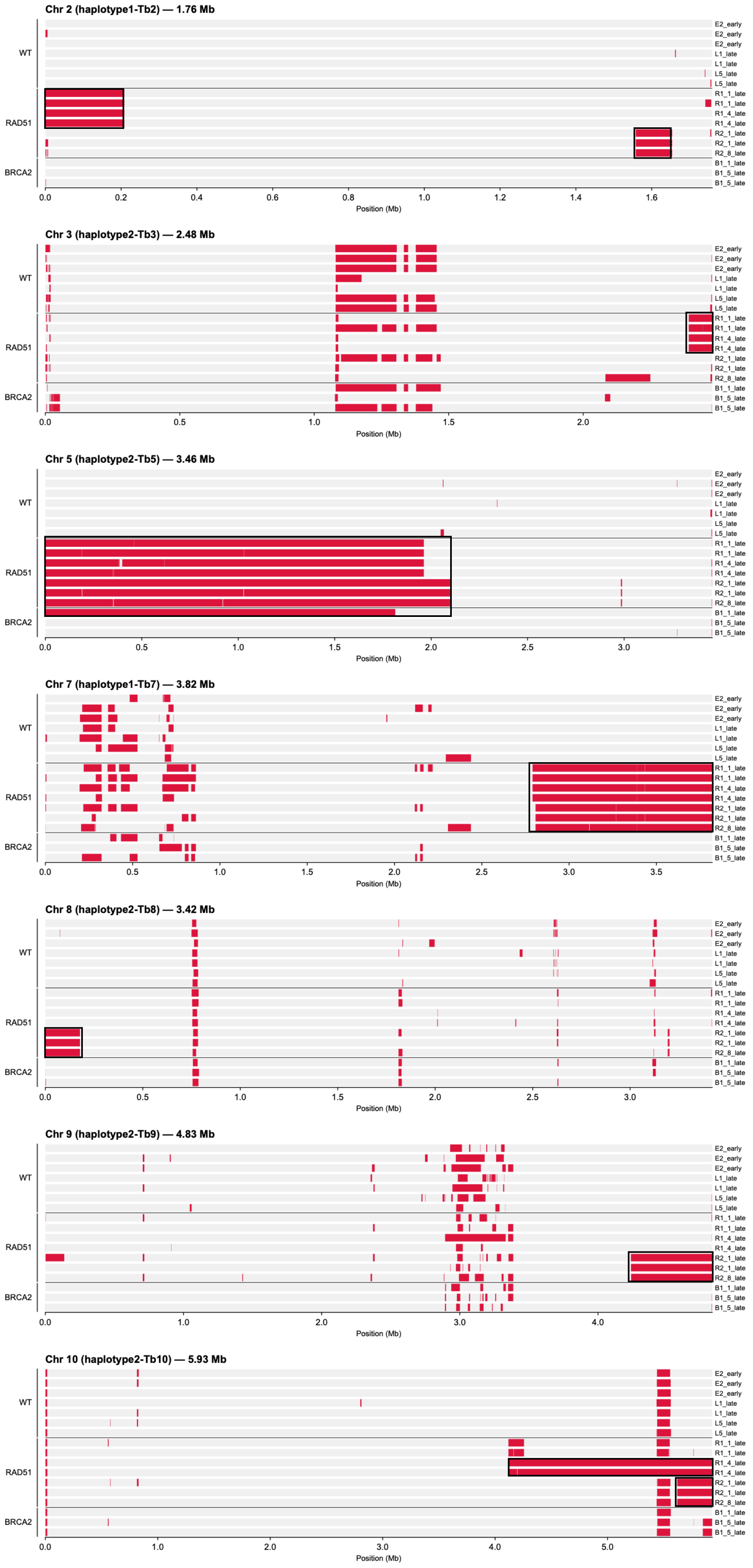
Chr 2 (haplotype1-Tb2) — 1.76 Mb
WT
RAD51
BRCA2
Position (Mb)
Chr 3 (haplotype2-Tb3) — 2.48 Mb
WT
RAD51
BRCA2
Position (Mb)
Chr 5 (haplotype2-Tb5) — 3.46 Mb
WT
RAD51
BRCA2
Position (Mb)
Chr 7 (haplotype1-Tb7) — 3.82 Mb
WT
RAD51
BRCA2
Position (Mb)
Chr 8 (haplotype2-Tb8) — 3.42 Mb
WT
RAD51
BRCA2
Position (Mb)
Chr 9 (haplotype2-Tb9) — 4.83 Mb
WT
RAD51
BRCA2
Position (Mb)
Chr 10 (haplotype2-Tb10) — 5.93 Mb
WT
RAD51
BRCA2
Position (Mb)

**Figure S14b**
